# Reliability of task-based fMRI in the dorsal horn of the human spinal cord

**DOI:** 10.1101/2023.12.22.572825

**Authors:** Alice Dabbagh, Ulrike Horn, Merve Kaptan, Toralf Mildner, Roland Müller, Jöran Lepsien, Nikolaus Weiskopf, Jonathan C.W. Brooks, Jürgen Finsterbusch, Falk Eippert

**Author notes:** **Address for correspondence:** Alice Dabbagh & Falk Eippert; Max Planck Research Group Pain Perception, Max Planck Institute for Human Cognitive and Brain Sciences, Stephanstraße 1a, 04103 Leipzig, Germany.

## Abstract

The application of functional magnetic resonance imaging (fMRI) to the human spinal cord is still a relatively small field of research and faces many challenges. Here we aimed to probe the limitations of task-based spinal fMRI at 3T by investigating the reliability of spinal cord blood oxygen level dependent (BOLD) responses to repeated nociceptive stimulation across two consecutive days in 40 healthy volunteers. We assessed the test-retest reliability of subjective ratings, autonomic responses, and spinal cord BOLD responses to short heat pain stimuli (1s duration) using the intraclass correlation coefficient (ICC). At the group level, we observed robust autonomic responses as well as spatially specific spinal cord BOLD responses at the expected location, but no spatial overlap in BOLD response patterns across days. While autonomic indicators of pain processing showed good-to-excellent reliability, both *β*-estimates and z-scores of task-related BOLD responses showed poor reliability across days in the target region (gray matter of the ipsilateral dorsal horn). When taking into account the sensitivity of gradient-echo echo planar imaging (GE-EPI) to draining vein signals by including the venous plexus in the analysis, we observed BOLD responses with fair reliability across days. Taken together, these results demonstrate that heat pain stimuli as short as one second are able to evoke a robust and spatially specific BOLD response, which is however strongly variable within participants across time, resulting in low reliability in the dorsal horn gray matter. Further improvements in data acquisition and analysis techniques are thus necessary before event-related spinal cord fMRI as used here can be reliably employed in longitudinal designs or clinical settings.

## 1. Introduction

Functional magnetic resonance imaging (fMRI) is a non-invasive method routinely used for brain imaging, with its first application in the human spinal cord about 30 years ago (Yoshizawa et al., 1996). Compared to the brain, the spinal cord is a more challenging environment for fMRI (Bosma & Stroman, 2014; Cohen-Adad, 2017; Eippert, Kong, Jenkinson, et al., 2017; Giove et al., 2004; Kinany, Pirondini, Micera, et al., 2022) and the number of studies using this technique has increased only slowly at first. However, the continued development and improvement of scanner hardware (Cohen-Adad et al., 2011; Lopez-Rios et al., 2023; Topfer et al., 2016), image acquisition protocols for spinal cord fMRI (Barry et al., 2021; Finsterbusch et al., 2013; Kinany,et al., 2022), shimming procedures (Finsterbusch et al., 2012; Islam et al., 2019; Tsivaka et al., 2023), denoising strategies (Brooks et al., 2008; Kong et al., 2012; Vannesjo et al., 2019) and software tools tailored to preprocessing and analyzing spinal cord data (De Leener et al., 2017, 2018; Rangaprakash & Barry, 2022) have made spinal fMRI more robust, sensitive and accessible, and accordingly has met with growing numbers of spinal fMRI studies more recently (Kinany et al., 2022; Landelle et al., 2021; Powers et al., 2018; Tinnermann et al., 2021).

Apart from a few notable exceptions (Conrad et al., 2018; Martucci et al., 2019, 2021; Rowald et al., 2022; Stroman, 2002; Stroman et al., 2004), most spinal cord fMRI studies have focused on cross-sectional designs in healthy volunteers, i.e. have not employed longitudinal designs or looked at the diagnostic or prognostic potential of spinal fMRI in clinical settings. This is different to brain imaging, the use of which in longitudinal settings and for biomarker development in the clinical context has been extensively discussed (Cole & Franke, 2017; Elliott et al., 2020; Khalili-Mahani et al., 2017; Kragel et al., 2021; Woo & Wager, 2016). Considering that the spinal cord is affected in a large range of neurological conditions – such as multiple sclerosis (Filippi & Rocca, 2013), neuropathic pain (Colloca et al., 2017) or spinal cord injury (Ahuja et al., 2017) – spinal cord fMRI could also be a valuable tool in clinical contexts, e.g., for tracking or predicting disease progression and treatment. However, the successful application of spinal cord fMRI in longitudinal settings and for diagnostic or prognostic purposes requires – at a minimum – achieving a high reliability of the method, with reliability being the extent to which measurement outcomes are consistent across different contexts. Test-retest reliability, for instance, describes the stability of a measure over time, i.e. it quantifies the precision of a method, or in other words, the expected variation over time, given that the underlying process of interest remains the same (Brandmaier et al., 2018; Lavrakas, 2008; Noble et al., 2021).

Studies assessing the test-retest reliability of spinal cord fMRI have mostly focused on resting-state signals (Barry et al., 2016; Hu et al., 2018; Kaptan et al., 2023; Kong et al., 2014; Kowalczyk et al., 2023; Liu et al., 2016; San Emeterio Nateras et al., 2016). Only three studies have examined task-related signal changes, with two of those using motor tasks (Bouwman et al., 2008; Weber et al., 2016b) and one using a pain task (Weber et al., 2016a). While these task-based studies provided important initial insights into the reliability of spinal cord fMRI, they had modest sample sizes (with at most N = 12) and mostly assessed reliability within a single scan session, thus circumventing some of the challenges inherent to longitudinal studies, such as repositioning of participants, and day-to-day variations in physiological state and mood (note that Bouwman and colleagues looked at a time-interval of 10 weeks, but only acquired data from three participants).

Here, we set out to provide a comprehensive assessment of the reliability of task-based spinal fMRI by investigating heat-pain evoked spinal cord BOLD responses. We chose the domain of pain for this endeavor for two reasons: on the one hand, changes in spinal cord processing are assumed to contribute to chronic pain (D’Mello & Dickenson, 2008; Kuner & Flor, 2017; Prescott et al., 2014) and on the other hand the development of pain biomarkers is currently a topic of intense interest (Davis et al., 2020; Leone et al., 2022; Mouraux & Iannetti, 2018; Sluka et al., 2023; Tracey, 2021), making spinal fMRI a prime candidate for inclusion in such clinical developments. In contrast to previous studies, we acquired data on two consecutive days using an identical experimental set up and a relatively large sample of 40 participants, as specified in an accompanying preregistration. We first analyzed the spatial distribution of the response across multiple spinal segments as well as its spread into the venous plexus surrounding the spinal cord. We then quantified the spatial consistency of the response patterns (using the Dice coefficient) and assessed test-retest reliability of BOLD responses in multiple ways (using the intraclass-correlation coefficient). Importantly, we simultaneously collected peripheral physiological data and compared their reliability to that of the BOLD data as only this allows for disambiguating between different causes for possibly low reliability in spinal cord BOLD responses, i.e. either poor data quality of spinal cord fMRI or variability in the underlying process (nociceptive processing).

## 2. Methods

### 2.1 Participants

40 healthy participants (20 female, mean age: 27.3 years, range: 18 – 35 years) participated in this study. This sample size was based on a preregistered power calculation (G*Power, Faul et al., 2007, version 3.1.9.7.), where we estimated that a sample of 36 participants would be necessary to detect a medium-sized effect (d = 0.5) with 90% power at an alpha-level of 0.05 when using a one-sample t-test against the baseline. All participants had normal or corrected-to-normal vision and a BMI < 24, were right-handed and gave written informed consent. The study was approved by the Ethics Committee of the Medical Faculty of the University of Leipzig.

### 2.2 Thermal stimulation

We employed phasic painful heat stimuli (duration: 1s, including 0.23s of ramp-up and ramp-down each, temperature: 48°C, baseline: 32°C), which were applied to the inner left forearm of the participants via an MRI compatible thermode with a ramp-speed of 70°C/s (surface of 9cm^2^, PATHWAY CHEPS; Medoc, Ramat Yishai, Israel). The stimuli were applied on five different areas of the inner left forearm (Figure 1), in order to minimize possible sensitization and habituation over runs.

**Figure 1.**
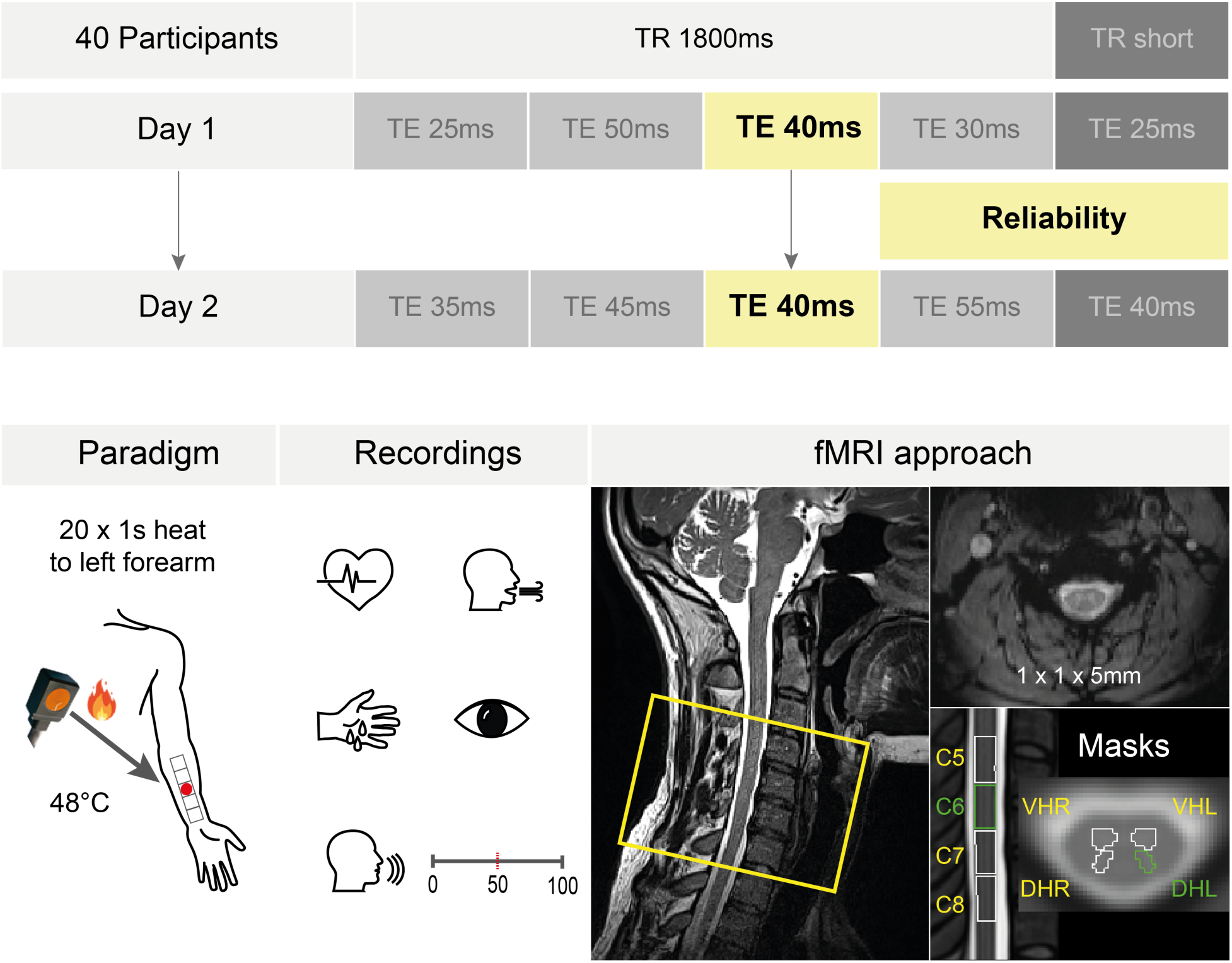
Experimental design. *Top*: 40 participants were measured on two consecutive days (sessions). On each day, we acquired five runs (each rectangle represents one run, also indicating the employed echo time [TE]). The runs were randomized across the two measurements days, with the exception that the Reliability Run (TE of 40ms, yellow rectangle) was measured on both days and is the focus of this report. Runs with a TR of 1800ms (medium grey rectangles) were combined with the Reliability Run for the Combined Runs average. Runs with a short TR (dark grey rectangles) were not included in this study. *Bottom*: Left: Before the measurements, we divided the area of the forearm into 5 equally sized patches, adapted to the individual proportions of the participant’s forearm. We drew the stimulation areas onto the left arm with a pen, to be able to target them easily when changing the thermode position in the scanner and to stimulate the same areas on the second day of the experiment. During each run, the participants received 20 heat stimuli with a duration of 1s and a temperature of 48°C. Across both days, we targeted the same skin patch for the Reliability Run. Middle: We measured heart rate, respiration, skin conductance and pupil dilation for each run and at the end of each run, participants were asked to rate the overall stimulus intensity using a numerical rating scale from 0-100 (0: no percept, 50: pain threshold, 100: unbearable pain). Right: Sagittal view of an example participant on the left, with the yellow rectangle indicating the rostro-caudal extent of the EPI slice stack (covering spinal cord segments C5 to C8, which translates to vertebrae C4 to C7). An axial example EPI slice (top right) demonstrates the data quality obtained with an in-plane resolution of 1×1mm (slice thickness: 5mm) and the masks employed are shown in the lower right, with the region of interest being the left dorsal horn in spinal cord segment C6 (depicted in green).

### 2.3 Experimental procedure

This study was part of a larger methodological project aimed at investigating the relationship between spinal cord BOLD responses and employed echo times. While we describe the entire data acquisition and experiment for the sake of transparency, here we solely focus on the data relevant for the issue of reliability – the echo-time dependence is the focus of an upcoming report.

At the beginning of the experiment, the participants were informed about the study and any remaining questions were discussed. We then outlined the five stimulation areas on the arm (most likely corresponding to dermatome C6 see Lee et al., 2008), using ink that remained visible over both days of the experiment. Before the main experiment started, the participants were familiarized with the heat stimulus by administering it twice to the right forearm, and then twice to each of the 5 possible stimulation areas on the left forearm. This served to minimize orienting / novelty responses, which could lead to detrimental movement at the beginning of a run.

After this familiarization, we prepared the participants for the placement in the scanner. We attached a breathing belt to measure respiration, as well as three electrodes to record the electrocardiogram (ECG; one electrode was placed on the left parasternal line at the level of the 1^st^ / 2^nd^ rib, another electrode on the left medioclavicular line at the level of the 9^th^ or 10^th^ rib, and the ground electrode on the left side of the chest, one hand-width below the armpit). Two electrodes were placed on the right hand to record skin conductance responses (SCR, one electrode on the thenar eminence, one electrode on the hypothenar eminence). The thermode was placed on the left arm. A custom-built MR-compatible extension mechanism (Mueller et al., 2024) was attached to the thermode that allowed for an easy repositioning of the thermode between scans from outside the scanner bore, without moving the participants and without changing the thermode pressure on the skin. After lying down on the scanner table, the participants were asked to tilt their head slightly towards their chest in order to minimize cervical lordosis (Cohen-Adad et al., 2021) and the isocenter was set approximately to the participants’ larynx. Before the experiment started, the eye tracker was calibrated to measure the pupil diameter throughout the experiment (eye tracker settings were validated or, if necessary, re-calibrated before the beginning of each run). The participants were instructed to avoid moving, avoid excessive swallowing, and to breathe normally (see Cohen-Adad et al., 2021) as well as to look at a fixation cross on a screen for the entire duration of each run while avoiding excessive blinking.

We split the experiment up across two consecutive days and measured five runs in each MRI session (Day 1 and Day 2). One run consisted of 20 trials, with one trial lasting between 11 and 13 seconds (1s stimulations and jittered inter-trial interval of 10-12s) and the duration of one run being four minutes and 48 seconds. We measured each run with a different echo time (TE; 7 runs: TE = 25ms, 30ms, 35ms, 40ms, 45ms, 50ms, 55ms) and a repetition time (TR) of 1800ms; two runs were additionally measured with the shortest possible TR (TE = 40ms & TR = 1560ms; TE = 25ms & TR = 1320ms). Splitting up the experiment across two days gave us the opportunity to assess the reliability of task-based spinal fMRI data. For this reason, we measured one run with a TE of 40ms (approximating T_2_* in the spinal cord, Barry & Smith, 2019) and a TR of 1800ms on each of the two measurement days (Fig. 1), which is the main focus of this study (from now referred to as the Reliability Run). In total, we therefore measured ten runs per participant, (nine TE&TR combinations and one additional Reliability Run), with five runs per day. Run order and targeted skin patch were pseudo-randomized and counter-balanced across participants, but kept identical across both days for the Reliability Run (see Fig. 1).

### 2.4 Data acquisition

The MRI data were acquired on a Siemens PRISMA FIT 3 Tesla scanner (Siemens, Erlangen, Germany), equipped with a whole-body radio-frequency (RF) transmit coil. We used a 64-channel RF head-and-neck coil, of which head coil element 7 and neck coil element groups 1 and 2 were utilized (all receive-only). We started the data acquisition with a localizer scan, followed by positioning the EPI slice stack and adjust volume (60 x 60 x 100 mm). A single EPI volume was then acquired, initializing the scanner’s “Advanced shim mode”, with the resulting shim applied to all following EPI acquisitions. The angle as well as centering of the adjust volume was identical to that of the EPI acquisitions, but it was slightly larger in superior-inferior-direction. We then acquired a z-shim reference scan that allowed for the automatic determination of the optimal z-shim moment for each slice of the EPI slice stack (Finsterbusch et al., 2012; Kaptan et al., 2022). A sagittal field map was obtained to estimate the static B0 field distribution. After this we measured a high-resolution T2-weighted structural scan for registration purposes, followed by two T2*-weighted ME-GRE scans to map T2* with two different resolutions. Finally, we measured the five functional runs. Prior to each functional run we acquired ten functional volumes with posterior-to-anterior phase encoding. In the following paragraph, we provide details on all protocols, except for the acquired field map, T2*-weighted ME-GRE and functional volumes measured with posterior-to-anterior phase encoding, since these were not utilized in the course of this study and will be described elsewhere (as will the EPI runs with shortened TR).

The EPI z-shim reference scan (TE: 40ms, total acquisition time: 55 seconds) consisted of 21 volumes with equidistant z-shim moments compensating for field inhomogeneities between +0.21 and −0.21 mT/m (in steps of 0.021 mT/m). The fMRI runs were measured via a single-shot 2D gradient-echo EPI sequence with 16 slices, covering the spinal cord from the 4^th^ cervical vertebra to the 1^st^ thoracic vertebra, with a resolution of 1 x 1 x 5mm (slice orientation: oblique axial; TR: 1.8s, TE different between runs: 25ms | 30m | 35ms | 40ms | 45ms | 50ms | 55ms, readout flip angle (FA): 75°, field of view (FOV): 128 x 128mm^2^, FOV position: centered rostro-caudally at level of 4th spinal disc, GRAPPA acceleration factor: 2, partial Fourier factor: 6/8, phase-encoding direction: AP, echo-spacing: 0.47ms, bandwidth per pixel: 1220 Hz/Pixel, slice angulation: perpendicular to each participant’s spinal cord, fat saturation and anterior and posterior saturation bands). All EPI acquisitions were performed with automatic slice-wise z-shimming (Finsterbusch et al., 2012; Kaptan et al., 2022). Three initial dummy shots were performed before the first functional image was acquired to achieve steady-state conditions. With the employed repetition time and flip angle, this approach brought all MR images to within 0.12% of the steady-state signal for gray matter, allowing us to include all images in the analysis. We also acquired a high-resolution T2-weighted structural scan via a SPACE sequence with a resolution of 0.8 x 0.8 x 0.8mm (slice orientation: sagittal, repetition time (TR): 1.5s, TE: 0.12s, FA: 120°, number of slices: 64, field-of-view (FOV): 256 x 256 mm^2^, GRAPPA acceleration factor: 3, bandwidth per pixel: 625 Hz/pixel; Cohen-Adad et al., 2021).

In addition to the MRI data, we acquired peripheral physiological data (respiration, heart rate, skin conductance and pupil diameter) throughout the entire experiment on both measurement days. Respiration, heartbeat, and skin conductance responses were recorded using a BrainAmp ExG system (Brain Products, Gilching, Germany) and pupil diameter was assessed via the Eyelink 1000 Plus system (SR research, Ottawa, Canada). Furthermore, after every run, the participants were asked to verbally rate the average intensity of the stimuli on a numerical rating scale (NRS) ranging from 0 to 100, where 0 translated to “no percept”, 50 marked the pain threshold and 100 translated to “unbearable pain”.

### 2.5 Peripheral physiological data analysis

#### 2.5.1 Heart period responses (HPR)

The ECG data were preprocessed using EEGLAB (Delorme & Makeig, 2004) and the FMRIB plug- in for EEGLAB, provided by the University of Oxford Centre for Functional MRI of the Brain (FMRIB) to remove MR-artifacts from the data traces recorded during functional runs (Niazy et al., 2005). Using in-house Matlab scripts, R-peaks were automatically detected, and manually corrected, if necessary. To obtain heart period time series, each inter-beat interval (IBI) was assigned to its following R-peak and the resulting IBI time series was linearly interpolated to achieve a sampling rate of 10Hz. Additionally, we filtered the IBI time series using a second-order Butterworth band-pass filter with cut-off frequencies at 0.01 Hz and 0.5 Hz (Paulus et al., 2016). The IBI traces were subdivided into event-epochs of −1 to 10s relative to stimulus onset and baseline-corrected to the average IBI within 5s before until stimulus onset. We then extracted the minimum of the IBI trace in an interval of 0 - 8s after stimulus onset of each trial and averaged the resulting 20 HPR values of the Reliability Run per day and participant. To test for differences in the HPRs between both days, we entered the average HPR of each participant and day into a pair-wise two-sided t-test.

#### 2.5.2 Skin conductance responses (SCR)

In two participants, SCR could not be recorded due to technical issues, leading to a sample size of 38 participants for SCR analyses. SCR data were down-sampled to 100Hz and low-pass filtered with a cut-off frequency of 1 Hz. The SCR traces were subdivided into event-epochs of −1 to 10s relative to stimulus onset and baseline-corrected to stimulus onset. We then extracted the peak of the skin conductance trace in an interval of 0 - 8s after stimulus onset of each trial and averaged the resulting 20 SCR values of the Reliability Run to acquire one average peak value per day and participant. To test for differences in the SCRs between both days, we entered the average SCR of each participant and day into a pair-wise two-sided t-test.

#### 2.5.3 Pupil dilation responses (PDR)

In six participants, eye tracking data of sufficient quality could not be recorded, leading to a sample size of 34 participants for pupil dilation analyses. Eyeblinks that were automatically detected by the EyeLink software were removed within a period of ± 100ms surrounding each blink. After the automatic blink detection, we manually corrected for any additional blinks or artifacts in the data trace by interpolating across the affected data segments. Blinking periods or otherwise missing data were interpolated linearly and the data were down-sampled to 100Hz and low-pass filtered with a cut-off frequency of 4 Hz. The pupil data traces were subdivided into event-epochs of −1 to 10s relative to stimulus onset and baseline-corrected to stimulus onset. We then extracted the peak of the pupil dilation trace in an interval of 0 – 4s after stimulus onset of each epoch and averaged the resulting 20 PDR values of the Reliability Run to acquire one average peak value per day and participant. To test for differences in the PDRs between both days, we entered the average PDR of each participant and day into a pair-wise two-sided t-test.

### 2.6 fMRI data analysis

Preprocessing and statistical analyses were carried out using FSL (version 6.0.3), SCT (Version 5.5) as well as custom MATLAB (version 2021a) and Python (version 3.9.13) scripts. The following procedures were carried out separately for the Reliability Run of each measurement day and participant.

#### 2.6.1 Preprocessing

##### Correction for thermal noise

As a first step, we applied non-local Marchenko-Pastur principal component analysis (MP-PCA, https://github.com/NYU-DiffusionMRI/mppca_denoise, Veraart et al., 2016) on the unprocessed EPI data of the Reliability Run to reduce thermal noise (Ades-Aron et al., 2021; Diao et al., 2021; Kaptan et al., 2023). The application of MP-PCA resulted in a substantial spinal cord tSNR increase (63.6%; from 11.69 before MP-PCA to 19.12 after MP-PCA), but only a marginal increase in spatial smoothness in the spinal cord (5.7%; from 1.23 before MP-PCA to 1.30 after MP-PCA; estimated via AFNI’s 3dFWHMx tool: https://afni.nimh.nih.gov/pub/dist/doc/program_help/3dFWHMx.html)

##### Motion correction

Motion correction was carried out in two steps. We first created a mean image of the 160 EPI volumes (after thermal noise correction). This mean image was used as a target image for the motion correction as well as to automatically segment the spinal cord. Based on the segmentation, we created a cylindrical mask, which was used to prevent adverse effects of non-spinal movement on the motion parameter estimation. Motion correction was then carried out slice-wise (allowing for x- and y-translations), using spline interpolation and a 2^nd^ degree polynomial function for regularization along the z-direction (De Leener et al., 2017). As a second step, we repeated the motion correction of the original time series of the Reliability Run, now using the mean image of the initially motion-corrected time-series as the target image.

##### Segmentation

In order to obtain a high-resolution segmentation of the spinal cord, we used the T2-weighted acquisition with 0.8mm isotropic voxels. For this purpose, we first applied the ANTs N4 bias field correction algorithm on the raw structural data to correct for intensity inhomogeneities due to RF coil imperfections (Tustison & Gee, 2010). As a next step, we denoised the structural data via Adaptive Optimized Nonlocal Means (AONLM) filtering (Manjón et al., 2010) to account for spatially varying noise levels and increase the SNR. To improve the robustness and quality of the final segmentation, we used an iterative procedure: the data were initially segmented using the SCT DeepSeg algorithm (Gros et al., 2019), smoothed along the z-direction using an anisotropic Gaussian kernel with 6mm sigma (in straightened space), and again segmented via the DeepSeg algorithm. To obtain a spinal cord segmentation of the EPI data, we used the mean image of the motion-corrected time series as input for SCT’s DeepSeg algorithm.

##### Registration to template space

While the statistical analyses on the individual level took place in each participant’s native space, group-level analyses were carried out in a common anatomical space defined by the PAM50 template of the spinal cord (De Leener et al., 2018). For the individual transformations from native to template space, we utilized the denoised and segmented structural T2-weighted image. In line with SCT’s recommended registration procedure (De Leener et al., 2017), the vertebral levels were identified and labeled, and the spinal cord was straightened. Using an iterative, slice-wise non-linear registration procedure based on segmentations, the structural image was then registered to the template. The resulting inverse warping field served to initialize the registration of the PAM50 template to the motion-corrected mean functional image via SCT’s multi-step non-rigid registration (De Leener et al., 2017). Based on this registration, we obtained a warping field to move the native-space mean functional image of each participant to template space using spline interpolation.

##### Correction for physiological noise

We employed several steps to reduce physiological noise. First, we identified volumes with excessive motion whose effects we aimed to remove during general linear model estimation. For this purpose, we calculated the root mean square difference between successive volumes (dVARS) as well as the root mean square intensity difference of each volume to a reference volume (refRMS) using FSL’s fsl_motion_outliers algorithm. Volumes presenting with dVARS or refRMS values three standard deviations above the time series mean were defined as outliers and individually modelled as regressors of no interest in subsequent analyses (on average 2% [range: 0.6% - 5.6%] of the 160 volumes per run were regarded as outliers). Second, the respiratory and preprocessed cardiac signals (see section 2.4.1 electrocardiogram) were used to create a physiological noise model (PNM), which approximates to what extent fMRI signal changes can be explained by respiratory and cardiac activity (Brooks et al., 2008). The approximation is based on the estimated cardiac and respiratory cycle phase during which the slices were obtained. The approach is derived from the retrospective image correction procedure (RETROICOR; Glover et al., 2000) and has been adapted for spinal cord fMRI to obtain slice-wise physiological regressors for subsequent analyses (Brooks et al., 2008; Kong et al., 2012). We extracted 32 noise regressors to estimate cardiac, respiratory and interaction effects as well as an additional regressor to model the cerebrospinal fluid (CSF) signal, which was derived from voxels in the CSF and spinal cord space that exhibited high levels of signal variance.

#### 2.6.2 Statistical analysis

##### General linear model

The statistical analysis of the fMRI data was based on the general linear model (GLM) approach implemented in FSL’s FEAT (FMRI Expert Analysis Tool; http://fsl.fmrib.ox.ac.uk/fsl/fslwiki/FEAT; Woolrich et al., 2001) and included spatial smoothing via FSL’s Susan tool with an isotropic 2mm (full width half maximum) Gaussian kernel as well as high-pass filtering at 100s. The first-level design matrix included a regressor for the heat stimulus onsets convolved with a double-gamma hemodynamic response function (HRF) as well as a temporal derivative. The following regressors of no interest were added to the design matrix for robust denoising: 33 slice-wise PNM regressors (describing cardiac, respiratory and CSF effects), two slice-wise motion regressors (describing movement along x and y; calculated during motion correction), and one regressor for each volume with excessive motion. From the first-level analysis we obtained a *β*-estimate map for the main effect of heat for each participant and day, which we registered to the template using the previously estimated warping fields.

##### Masks

For the subsequent analyses, we used multiple masks. In template space, we first derived z-coordinates to divide the spinal cord according to the spinal segmental levels C5 to C8 (which are the levels covered by our EPI slice-stack). The coordinates for each segment were obtained from SCT (version 6.1, De Leener et al., 2018) and are based on findings from Frostell et al., (2016). Segmental masks for the four gray matter horns of the spinal cord were derived by cropping the unthresholded probabilistic gray matter masks of each gray matter horn according to the same segmental coordinates. Additionally, we utilized segmental masks for the four quadrants of the cord, encompassing both gray matter and white matter, using the same segmental coordinates. The mask of the left dorsal horn in segment C6 (number of isotropic 0.5mm voxels: 502) was used to investigate the main effect of heat, as well as the reliability between the days, along with the mask of the right ventral horn of segment C6 (number of isotropic 0.5mm voxels: 719), which was defined as a control region. To assess BOLD activity patterns beyond the target region, we used a cord mask of spinal segment C6 dilated by 6 voxels (i.e., including an area occupied by draining veins), which we then subdivided into 4 quadrants (left dorsal, number of isotropic 0.5mm voxels: 5888; left ventral, number of isotropic 0.5mm voxels: 6671; right dorsal, number of isotropic 0.5mm voxels: 5888, right ventral, number of isotropic 0.5mm voxels: 6671).

##### Average and day-wise BOLD responses

To first investigate whether phasic heat stimulation as employed here evokes a significant BOLD response at all, we averaged the normalized *β*-maps over both days within each participant and then submitted the resulting 40 *β*-maps to a one-sample t-test. Correction for multiple comparison was carried out via voxel-wise non-parametric permutation testing as implemented in FSL’s randomise algorithm (Winkler et al., 2014) in an anatomically informed target region (left dorsal horn, segment C6) at a threshold of p_FWE_ < 0.05 (family-wise error corrected). Second, we also aimed to test for significant responses on each of the two days using exactly the same statistical procedure, but now using the 40 *β*-maps from each day as input. Finally, we aimed to test for an overlap of significant responses across both days (i.e., a conjunction) and therefore created a binary mask of the significant voxels of each day, which we subsequently multiplied with each other to determine significant voxels overlapping across both days.

#### Spatial specificity

We also aimed to describe the spatial organization of the BOLD responses in the part of the spinal cord covered by our slice-stack. For this purpose, we performed a one-sample t-test (again within a permutation-testing framework) using the averaged *β*-estimates over both days within a cord mask including spinal segments C5 to C8. From the resulting uncorrected group-level p-maps, we assessed the number of voxels surviving liberal thresholding at p < 0.001 uncorrected in each spinal segment. We then calculated what percentage of the total number of active voxels in the respective cord segment were located in the dorsal left, dorsal right, ventral left and ventral right cord quadrant (including both gray and white matter parts of the cord). In order to supplement this analysis with more fine-grained information regarding gray matter responses, we additionally assessed the number of supra-threshold voxels within the four gray matter horns.

Finally, we aimed to assess to what degree BOLD responses also occur ‘outside’ the spinal cord proper. While our target tissue of interest is the gray matter of the dorsal horn, this is drained through a hierarchy of veins: small veins coalesce into radially-oriented intramedullary veins, which further drain into the circumferential spinal veins of the superficial pial venous plexus, a structure itself permeated by longitudinal veins. From here, the blood drains into the internal vertebral plexus before progressing to the external vertebral plexus, ultimately joining the systemic circulation (Thron, 2016). Taking into account this network of draining veins outside the gray matter is crucial for spinal cord fMRI, as these venous pathways can influence the spatial specificity of the BOLD response, potentially diluting the signal across a larger area than the region of neuronal activity. To better understand the resulting signal spread, we performed an additional group analysis within an extended region encompassing spinal segment C6, dilated by 4 voxels (i.e., 2mm, covering parts of the venous drainage system, such as the pial venous plexus) on a slice-by-slice basis.

### 2.7 Reliability

#### 2.7.1 Contextual differences

Before quantifying the reliability of the response measures, we tested for differences between the scanning days that could explain changes in the BOLD fMRI results, such as differences in the general physiological state of the participants and in the fMRI data quality. To assess differences in the physiological state of participants, we calculated three metrics: heart rate, heart rate variability and spontaneous fluctuations of electrodermal activity. All three metrics served to describe underlying differences in tonic autonomous nervous system activity, e.g. due to stress or emotional arousal (Bach et al., 2010; Berntson et al., 2017; Dawson et al., 2017). Heart rate was quantified as beats per minute (bpm) and heart rate variability (HRV) was calculated as the root mean square of successive peak-to-peak interval differences between normal heartbeats (RMSSD) in milliseconds. Spontaneous fluctuations in electrodermal activity were calculated by i) setting up a GLM where stimulus onsets were convolved with a canonical skin conductance response function (implemented in PsPM: https://bachlab.github.io/PsPM/; Bach et al., 2009, 2013) and ii) then using the residual activity (i.e. after removal of modelled stimulus-evoked responses) to calculate the area under the curve of the remaining skin conductance traces (Bach et al., 2010). To describe the overall quality of the fMRI data, we i) estimated motion by calculating the root mean square intensity differences of each volume to a reference volume and ii) calculated the temporal signal-to-noise ratio (tSNR) of the motion-corrected EPI data.

#### 2.7.2 Intra-class correlation coefficient

To assess the reliability of responses to painful heat stimulation across two days, we calculated the intra-class correlation coefficient according to Shrout & Fleiss (1979), a widely used statistical measure to assess the reliability of repeated measurements. Specifically, ICC(3,1) serves to assess the consistency of measurements across different occasions or days, and is defined as the ratio of the between-participant variance and the total variance (Caceres et al., 2009). ICC(3,1) was calculated using the following formula:

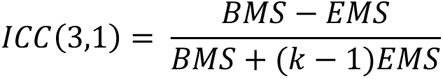

The formula includes three components: BMS (Between Participants Mean Squares), which represents the variance between different participants; EMS (Error Mean Squares), representing the residual variability, which includes inconsistencies in repeated measurements for the same participant; and k, indicating the number of measurements (Caceres et al., 2009). The numerator (BMS - EMS) reflects the variability between participants after accounting for measurement errors, while the denominator (BMS + (k - 1) * EMS) combines the total variability between participants with measurement errors adjusted by the number of measurements. This ratio quantifies the consistency of the measurements and indicates the degree to which participants maintain their relative ranking across days (Caceres et al., 2009; Liljequist et al., 2019). The calculation of the reliability coefficients for all of the measures described below was implemented via the Python package Pingouin (version 0.5.3, Vallat, 2018). The interpretation of the resulting reliability estimates followed the conventions by Cicchetti (1994), where ICC values smaller than 0.4 indicate poor reliability, values from 0.4 - 0.59 imply fair reliability, values from 0.6 - 0.74 represent good reliability and values from 0.75 - 1.0 are defined as excellent reliability.

#### 2.7.3 Subjective and peripheral physiological responses

We calculated the test-retest reliability for the verbal ratings of pain intensity (data from two participants were missing due to technical issues, resulting in N = 38) as well as for SCR, PDR, and HPR. For verbal ratings, we employed the single rating obtained after all trials and for the peripheral physiological measures, we employed the peak response value (averaged across trials) of each participant obtained on each of the two days (see sections 2.5.1., 2.5.2, 2.5.3 for closer description of peak value extraction).

### 2.7.4 BOLD responses

To assess the test-retest reliability of spinal cord heat-evoked BOLD responses across two consecutive days, we extracted *β*-estimates for the main effect of heat for each participant and session. We assessed the reliability through the application of different anatomical masks (see section 2.6.2, under *Masks*). Two masks covered the area of interest, one of them limited to the gray matter horn, while the other mask incorporated draining vein territory, and two further masks captured the gray matter in a control region as well the draining vein territory adjacent to the control region. The first mask was the left dorsal horn mask on spinal segment C6, the second mask was the left dorsal quadrant of the dilated cord mask on the level of C6, the third mask was the right ventral horn on C6 and the fourth mask was the right ventral quadrant of the dilated cord mask of C6. The calculation of reliability in the left dorsal horn, was based on three metrics, namely the mean *β* over the entire region of interest, the peak *β* estimate in the ROI regardless of its location within the ROI, and finally the average of the top 10% *β* values in the ROI. We also calculated the reliability for the ROI mean, peak value and average of the top10% using the z-scores from the z-maps, since the z-scores scale the parameter estimate (*β* maps) by the standard error of each voxel, thereby considering the underlying variation within runs.

The reliability assessment described so far aimed to quantify the similarity of the response amplitudes, quantified via the *β* estimates or z-scores, over both days. However, in the context of fMRI, not only the response amplitude holds importance but also the spatial patterns of the response – specifically, we wanted to know whether the BOLD response on Day 1 occurred in the same location as the BOLD response on Day 2. To compare the spatial patterns of the BOLD responses between days, we calculated Dice coefficients, which quantified the amount of overlap of the active voxels in the left dorsal horn in spinal segment C6 (Rombouts et al., 1999; Wilson et al., 2017).

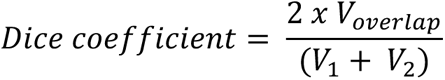

V_1_ and V_2_ define the number of active, i.e., above-threshold voxels on each day, and V_overlap_ is the number of voxels that overlap. We calculated Dice coefficients on the group and individual level using binarized statistical maps. On the group level we binarized the uncorrected p-maps at the thresholds 0.001, 0.01 and 0.5, and on the individual level we binarized the z statistic image for the main effect of heat, thresholded at +/-1.96 (i.e., p < 0.05 uncorrected).

## 2.8 Post-hoc analyses

As reported in the Results section, test-retest reliability was low for the peak activation in the left dorsal horn on C6. Additionally, in the same ROI there was no spatial overlap between the group-level results of both days, accompanied by a low Dice coefficient for even rather liberal thresholds. To investigate possible reasons for this surprising lack of response consistency, we carried out three further sets of analyses, which we had not specified in the preregistration.

### 2.8.1 Increasing the number of runs

The Reliability Run was optimized for assessing reliability in terms of keeping the measurement parameters and stimulation position on the arm identical across days. However, since one run consisted only of 20 trials and our stimulus duration of 1s was relatively short, this data is likely noisier than data of spinal fMRI paradigms with more trials or more prolonged stimuli, which might explain the low reliability. For this reason, we also investigated the fMRI activation maps and reliability metrics using the average over multiple runs per day, resulting in a four-fold increase of trials, since we included all runs with a TR of 1800ms (including the Reliability Run), i.e. four runs per day (the two additional runs with shorter TRs were excluded from this analysis due to the different TR and flip angle employed in those acquisitions).

The preprocessing followed the steps described above and was done separately for each day. To bring all individual runs into a common space, motion correction was carried out exactly as described in section 2.6.1, however, instead of correcting a single run we concatenated all suitable runs of the respective day and motion-corrected the entire concatenated time series. Registration to template space followed the identical procedures as above, only now the mean image of the concatenated and motion-corrected time series served as the destination image. For all subsequent analyses we used the same procedures as described above (section 2.6.2), with the difference that the *β* maps of the individual runs were averaged across runs and only then entered the group-level analysis. For the rating and all peripheral physiological data, we also combined the data of the four runs per day and calculated the reliability coefficients accordingly. The results of this analysis are referred to as “Combined Runs”.

### 2.8.2 Accounting for spontaneous activations

Another cause of the low reliability of task-evoked BOLD responses could be spontaneous fluctuations in the BOLD signal, which were not accounted for in the GLM, and which might increase trial-to-trial variability. A study by Fox and colleagues (2006) showed that trial-to-trial BOLD response variability in the left somatomotor cortex could be reduced by regressing out the BOLD signal of the right somatomotor cortex. The authors argue that the spontaneous fluctuations of both regions correlated due to the interhemispheric connectivity between the regions. Regressing out the signal of the opposite hemisphere mainly decreased noise, whereas the accompanied reduction of the task-relevant signal was non-significant. Since previous spinal fMRI studies have found evidence for resting-state functional connectivity between the left and right dorsal horn (Barry et al., 2014, 2016; Eippert, Kong, Winkler, et al., 2017; Harita & Stroman, 2017; Kaptan et al., 2023; Kinany et al., 2020; Kong et al., 2014; Vahdat et al., 2020), we aimed to test if a similar analysis strategy could decrease noise due to spontaneous fluctuations, and improve reliability (though we are aware that this could also be negatively affected e.g. due to pain-induced responses in contralateral dorsal horns (Fitzgerald, 1982). For this purpose, we extracted the time series of the contralateral (right) dorsal horn of each slice, and used it as an additional slice-wise regressor in the GLM to regress out spontaneous fluctuations in the ipsilateral (left) dorsal horn. Otherwise, the analysis followed the procedure outlined above.

### 2.8.3 Within-run reliability

Given the low reliability across days, we wanted to assess if reliability would be equally low within runs, as such comparisons would not involve the potentially detrimental impact of repositioning the participants in the scanner as well as possibly imperfect matches of the normalized parameter estimate maps in template space. For this purpose, we adopted a split-half approach and divided the Reliability Run into two subsets: odd and even trials for the odd-even reliability analysis, or the first and second half for the early-late reliability analysis, with the corresponding trial regressor entered in the general linear model (GLM). Subsequently, we obtained two *β* maps from both the odd-even and early-late GLMs, representing the respective trial selections. These *β* maps were then subjected to the spatial normalization procedure described in section 2.6.1. Reliability coefficients (ROI mean, ROI peak, average of top 10% *β* estimates extracted for each participant) were calculated between the respective trial selections of both approaches, separately for each day and the resulting ICC scores were averaged across days. We also calculated both within-run reliability measures for SCR, PDR and HPR, calculating the respective response peaks for each set of trials and averaging ICC values across days.

### 2.8.4 Correlations between BOLD and non-BOLD response measures

In an additional exploratory analysis inspired by a reviewer’s comment, we assessed the across-participant correlations between peripheral physiological as well as subjective responses and BOLD responses (results are reported in Supplementary Table 3). For this purpose, for every participant we averaged the top 10% *β* estimates and z-scores in the left dorsal horn of C6, along with SCR, PDR, HPR and subjective ratings across days. We then calculated Pearson’s r for each of the eight correlations (only including participants with responses in both variables): *β* estimates with SCR, HPR, PDR, and rating as well as z-scores with SCR, HPR, PDR, and rating. Since we assumed that higher BOLD responses would go along with stronger peripheral physiological and subjective responses (positive associations expected for all responses but HPR, as here a negative-going response indicates cardiac acceleration as is typical in response to nociceptive input), we base our results on one-tailed p-values as indicators of statistical significance of the correlation strength.

### 2.8.5 Correlations between BOLD parameter estimates and indicators of data quality

Inspired by a reviewer’s comments, we calculated correlations between changes in data quality metrics and BOLD parameter estimates across days, as this should allow for insights into possible data quality contributions to across-day reliability of BOLD responses. For every participant we calculated the difference from Day 1 to Day 2 for i) motion estimates, ii) normalization quality estimates and iii) indicators of participant positioning. Motion estimates were obtained via root mean square intensity differences of each motion-corrected volume to reference volume (see refRMS, section 2.6.1). Normalization quality was estimated via computing Dice coefficients between the segmentation of the normalized EPI and the PAM50 cord mask (see section 2.7.4 for Dice coefficient). Participant positioning estimates were obtained by calculating the angulation of the slice stack relative to the direction of the B0 field, since the slice stack was always positioned to be orthogonal to the longitudinal axis of the spinal cord (see Fig. 1 for example). The angle between the normal vector of the slice package extracted from the DICOM header and the scanner’s z-axis (0,0,1) therefore serves as a proxy for the positioning of the neck – and thus the orientation of the draining veins – relative to B0. For each of these measures, we correlated the absolute difference across days with the absolute difference of BOLD parameter estimates across days, quantified as the top 10% *β* estimates and z-scores of the left dorsal horn in spinal cord segment C6. Since we expected a positive correlation (i.e., greater differences across days would be associated with greater variation of BOLD responses), we report one-tailed p-values alongside the correlations in Supplementary Table 4.

### 2.9 Open Science

This study was preregistered before the start of data acquisition and the preregistration is openly available on the Open Science Framework (https://osf.io/a58h9); differences between the analyses suggested in the preregistration and the analyses carried out here (as well as the reasons behind these changes) are listed in the Supplementary Material. The underlying data and code are currently only accessible to reviewers, but will be made openly available upon publication via OpenNeuro and GitHub, respectively. The intended data-sharing via OpenNeuro was mentioned in the Informed Consent Form signed by the participants and approved by the Ethics Committee at the Medical Faculty of the University of Leipzig.

## 3. Results

### 3.1 Behavioral and physiological responses

Across both days, the participants reported an average stimulus intensity of 71.7 (Fig. 2, left, n = 38, SD = 12.1), indicating that the employed heat stimuli were perceived as clearly painful (responses greater than 50 indicated pain). This subjective percept was accompanied by robust physiological changes (Fig. 2), as evidenced in skin conductance responses (SCR), pupil dilation responses (PDR) and heart period responses (HPR). All measures showed rather similar responses when compared across days (t_rating_(37) = 0.15, p_rating_ = 0.88; t_SCR_(37) = 0.86, p_SCR_ = 0.39; t_PDR_(33) = 0.67, p_PDR_ = 0.51; t_HPR_(39) = 0.48, p_HPR_ = 0.63; Fig. 2).

**Figure 2.**
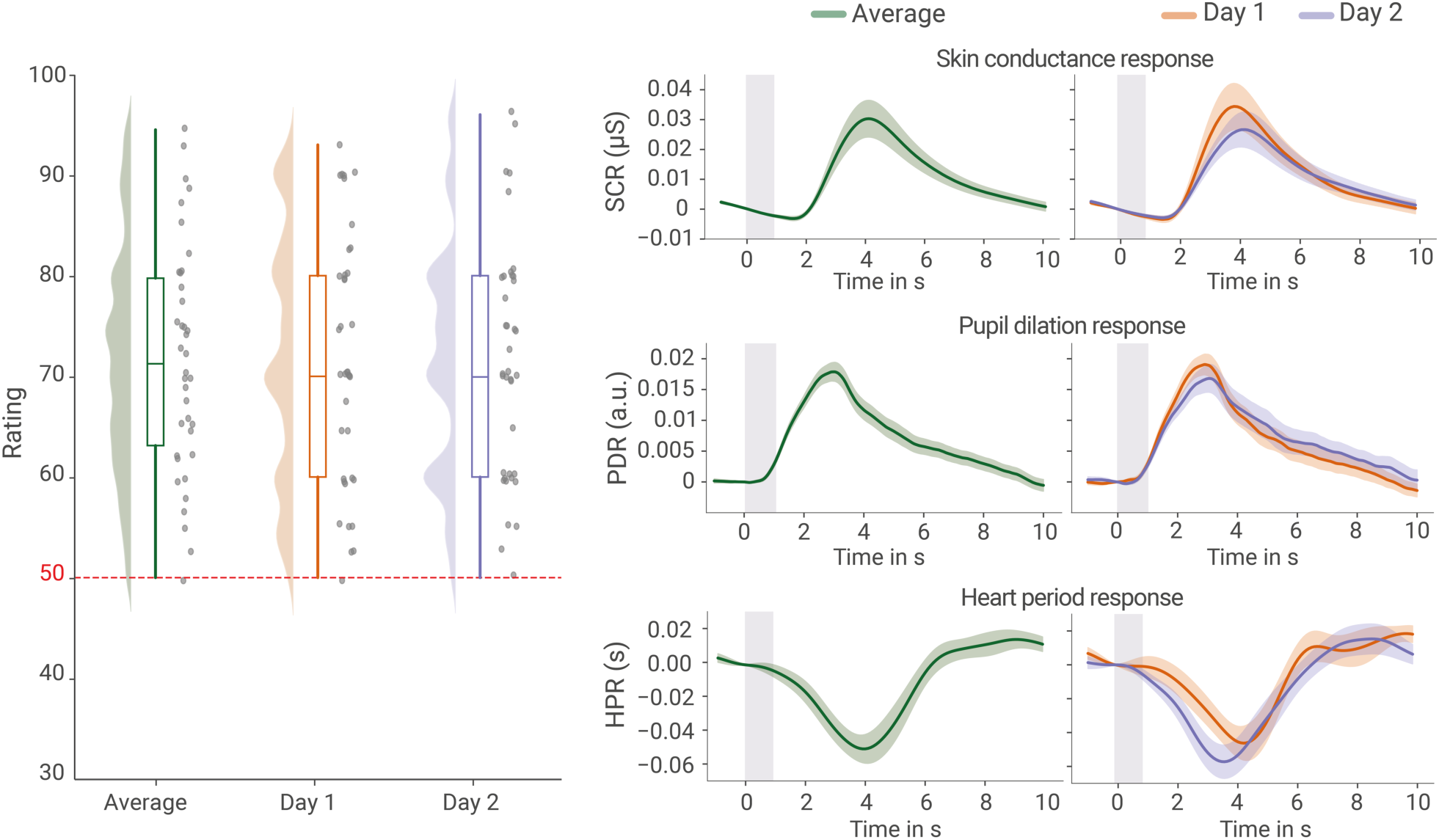
Subjective and peripheral physiological responses. Left: verbal ratings of stimulus intensity on a numerical rating scale with 50 indicating the pain threshold; half-violins and boxplots depict the distribution over participants and grey dots show the raw values (jittered slightly for visualization purposes). Right: Averaged traces of the skin conductance response (SCR), pupil dilation response (PDR) and heart period response (HPR) in response to the stimulus, with error bands reflecting the standard error of the mean across the group and the gray rectangle representing the stimulus duration.

### 3.2 BOLD responses: amplitudes

Averaged across days, we observed a significant response in the ipsilateral dorsal horn in spinal segment C6 (t(39) = 4.51, p_corr_ = 0.002, 61 supra-threshold voxels; Fig. 3). Using the same analysis parameters, we also observed significant responses for each day separately (t_day1_(39) = 3.58, p_corr_ = 0.035, 2 supra-threshold voxels; t_day2_(39) = 4.50, p_corr_ = 0.0018, 48 supra-threshold voxels). When comparing the spatial pattern of active voxels (at a threshold of p < 0.05 corrected) for Day 1 and Day 2, there was no overlap, with the active voxels of Day 1 being located consistently more caudal in segment C6 compared to the active voxels of Day 2 (Fig. 3, sagittal view), despite the heat stimulation occurring at the identical location on the forearm.

**Figure 3.**
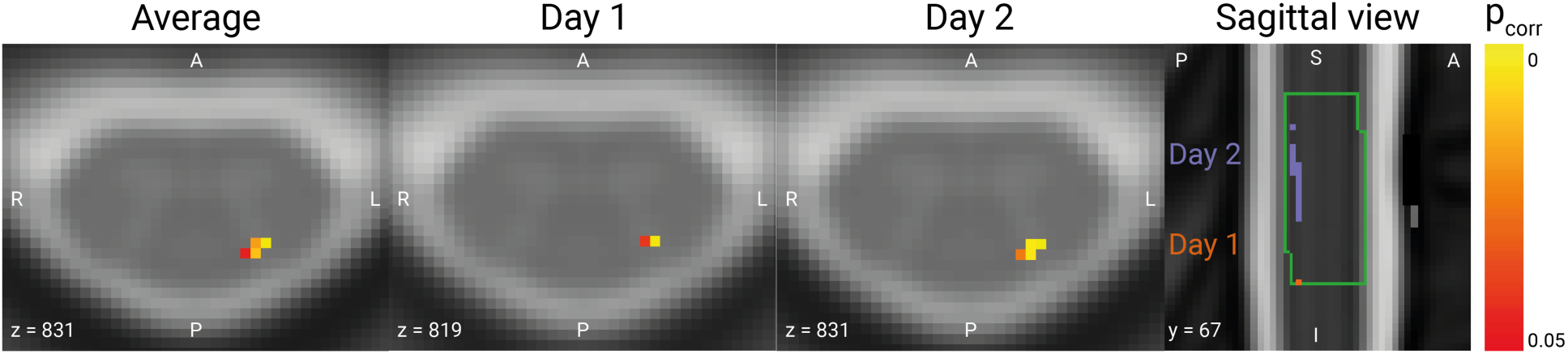
BOLD responses. Axial view of group-level results in the left dorsal horn in spinal segment C6 thresholded at p_FWE_ < 0.05, with the average across both days depicted on the very left, followed by Day 1 and Day 2. The rightmost plot shows a sagittal view of the activation maps of both days, with purple voxels belong to Day 1 while red voxels belong to Day 2; the green outline marks spinal cord segment C6 for visualization purposes. Data are overlaid on the T2*-weighted PAM50 template in axial views, and on the T2-weighted PAM50 template in the sagittal view.

### 3.3 BOLD responses: spatial specificity

#### 3.3.1 Entire spinal cord

To assess the spatial specificity of BOLD responses, we used the group-level results of the across-day average within the cord mask from spinal segment C5 to C8. We counted the number of active voxels in each segment using a liberal threshold (p < 0.001 uncorrected) and assessed what percentage of those voxels was located in each of the 4 cord quadrants: left dorsal, right dorsal, left ventral and right ventral (Fig. 4; for exact percentages and day-wise results see Supplementary Table 1). The highest number of voxels that survived thresholding was located in spinal cord segment C5, followed by C6 and C7, with C8 holding the lowest number of supra-threshold voxels. As can be seen in the percentages, segments C6 demonstrated the highest level of spatial (i.e., neuroanatomical) specificity, followed closely by C5. In both segments, the majority of active voxels were concentrated in the left dorsal quadrant, and a smaller number of active voxels were found in the right dorsal quadrant, with a relatively small percentage of active voxels observed in the ventral region. Conversely, the above-threshold voxels in C7 were mostly located in the right ventral quadrant, and in spinal segment C8 in the right dorsal part.

**Figure 4.**
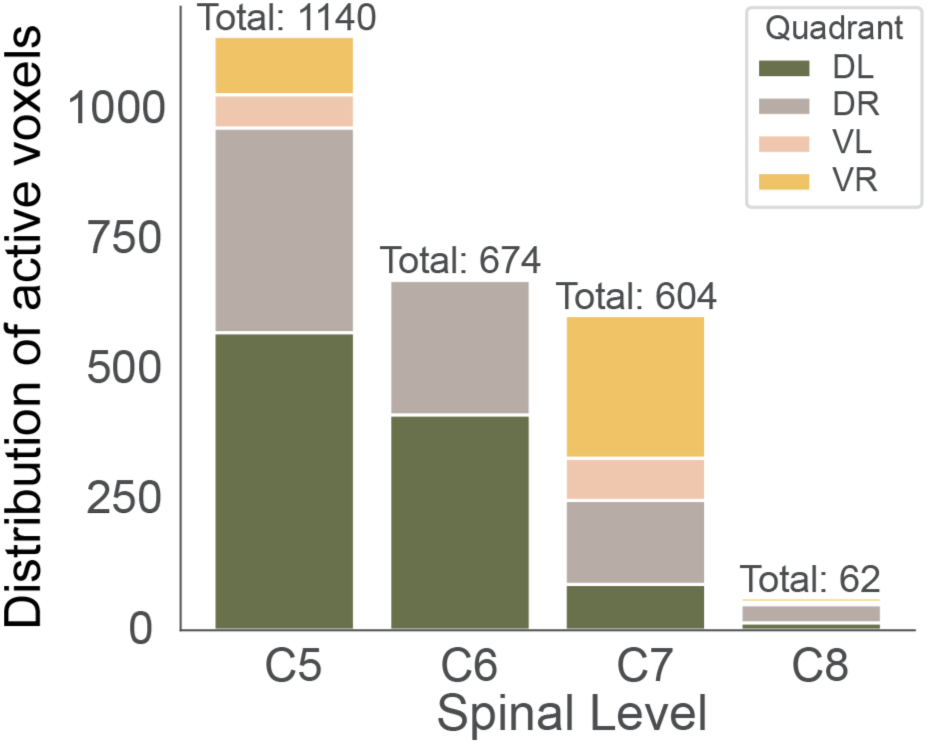
Spatial specificity of BOLD responses across cord quadrants. Number of supra-threshold voxels across the four cord quadrants of all spinal segments from C5 to C8. The total on top of each bar shows the overall number of active voxels in the entire cord mask of each level. Colors indicate the number of active voxels across the dorsal left (DL) and right (DR) as well as ventral left (VL) and right (VR) quadrants. All results shown here are based on the group-level across-day average (uncorrected p < 0.001).

#### 3.3.2 Gray matter

In order to obtain a more detailed understanding of spatial specificity in the spinal cord gray matter (instead of the cord quadrants, as reported above), we also projected all active voxels in the four gray matter horns onto an exemplary spinal cord slice, either from all segments (grey dots in Fig. 5A,) or only from target segment C6 (red dots in Fig. 5A) and furthermore visualized their distribution along the left-right and dorsal ventral axis. Across all segments, the highest number of active voxels was located in the left dorsal horn, but a substantial number of active voxels was also present in the other horns. Conversely, for our target segment C6, the clear majority of voxels is located in the ipsilateral dorsal horn, with a lesser number of voxels in the contralateral dorsal horn and no active voxels in the ventral horns.

**Figure 5.**
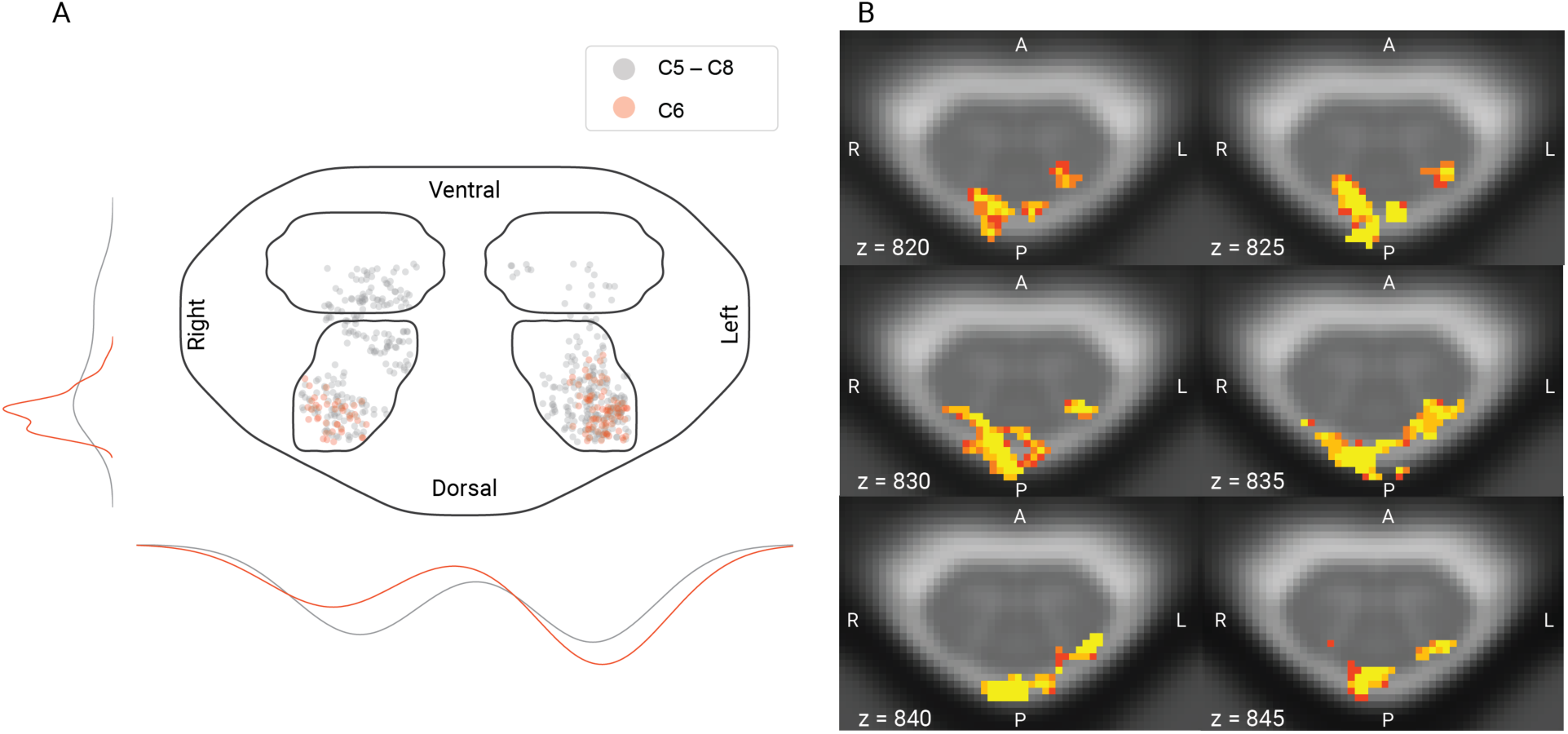
Spatial specificity of BOLD responses. A: Positions of supra-threshold voxels in the gray matter horns of all spinal segments from C5 to C8, collapsed over z (grey), vs. only in spinal segment C6 (red), with jitter added for visualization purposes. The lines on outside of plot show the distribution of the voxels across hemicords (lines on the left average over the left and right horns, lines on the bottom average over the dorsal and ventral horns); colors indicate the employed mask (red: only spinal segment C6, grey: collapsed over z). The mask used for visualization here was obtained by combining a slice of the cord and all four GM horn masks in C6. B: Six example slices across segment C6 showing supra-threshold voxels within a dilated cord mask to allow for observing draining vein responses. All results shown here are based on the group-level across-day average (uncorrected p < 0.001).

#### 3.3.3 Surrounding tissue

To investigate the impact of draining veins on the location of BOLD responses, we also assessed the spatial pattern of active voxels (p < 0.001 uncorrected, across-day group-level average) in a mask of spinal segment C6 that also included the venous plexus (Fig. 5B). Several aspects are worth noting here. First, in line with the previously presented data, there is almost no ventral horn activation and thus also no BOLD responses in the venous plexus on the anterior surface of the cord. Second, gray matter responses are consistently present throughout segment C6 in the ipsilateral dorsal horn and with lesser prominence also in the contralateral dorsal horn. Most importantly though, the strongest BOLD responses are actually observed at the dorsal surface of the cord in the region of the veins draining the dorsal cord: these responses are evident both ipsi- and contralaterally, at times spanning both the left and right dorsal surface.

### 3.4 Reliability

#### 3.4.1 Physiological state and data quality across days

To test for differences in the participants’ general physiological state across days, we calculated run-wise heart rate, heart rate variability, and spontaneous fluctuations of the electrodermal activity (Fig. 6A-C). While heart-rate showed a slight increase from Day 1 to Day 2 (t(39) = −2.15, p = 0.04), neither heart rate variability (t(39) = 1.23, p = 0.22) nor spontaneous fluctuations in electrodermal activity (t(37)=0.04, p = 0.97) showed significant differences across days. To assess fMRI data quality across days, we investigated motion effects (quantified as root mean square intensity difference to a reference volume for each run per participant and day; Fig. 6D) and temporal signal-to-noise ratio (tSNR; Fig. 5E) after motion correction. Neither motion effects (t(39) = 1.1, p = 0.28) nor tSNR (F(1,38) = 0.3, p = 0.59) showed significant differences across days; for tSNR this pattern held across all slices.

**Figure 6.**
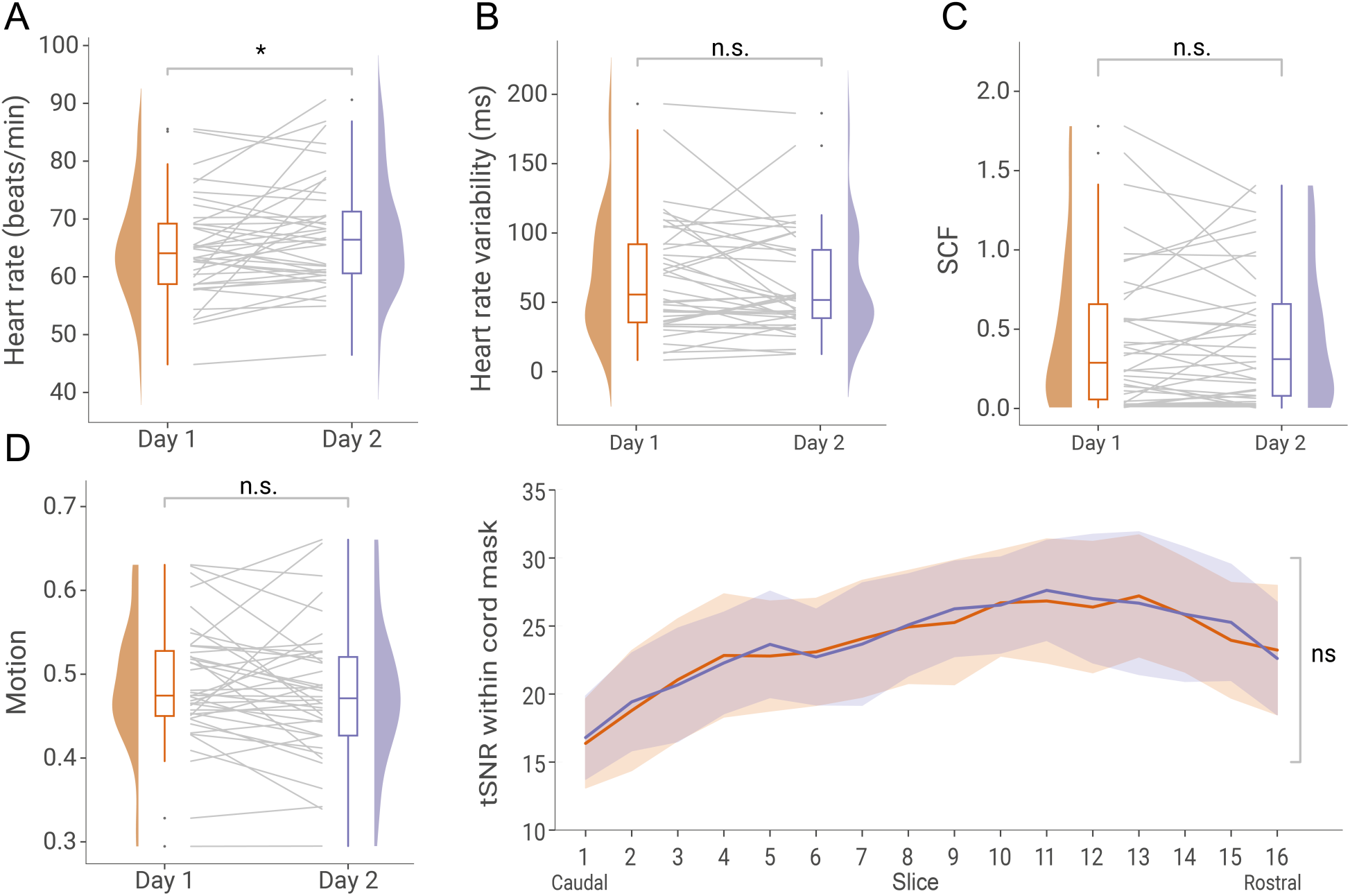
Physiological state and data quality across days. A – D show the average physiological state or fMRI quality indicators of each participant and day, visualized via box-plots, half-violin plots and grey lines that indicate participant-wise changes across days. A) Heart-rate quantified as beats per minute. B) Heart-rate variability quantified as root mean square of successive differences between normal heartbeats in ms. C) Spontaneous fluctuations in skin conductance (SCF) quantified as area under the curve. D) fMRI motion quantified as root mean square intensity differences of each volume to reference volume. E) fMRI signal quality quantified as temporal signal-to-noise ratio (tSNR) within a cord mask of each slice.

#### 3.4.2 Behavioral and peripheral physiological test-retest reliability

As a positive control analysis, we first assessed the reliability of the behavioral and peripheral physiological measures in order to ascertain that responses to noxious thermal stimulation can in principle be reliably assessed (Fig. 7, Table 1; Supplementary Figure 1). Subjective ratings (ICC = 0.72), skin conductance (ICC_SCR_ = 0.77) and heart period (ICC_HPR_ = 0.77) exhibited good-to-excellent test-retest reliability, whereas pupil dilation (ICC_PDR_ = 0.34) showed poor test-retest reliability.

**Figure 7.**
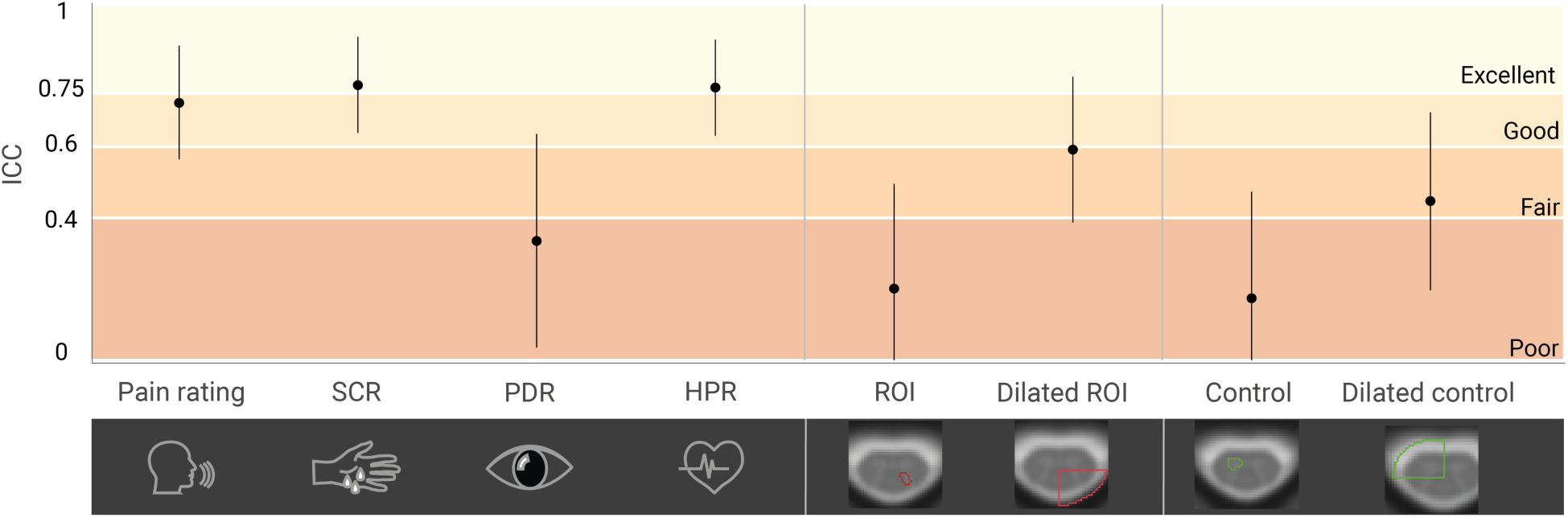
Test-retest reliability across both days for subjective ratings, peripheral physiological data and BOLD response amplitudes. Reliability is indicated via ICCs (plotted as dots with 95% CI represented as a line). ICCs are reported for (from left to right) verbal ratings, skin conductance response amplitude (SCR), pupil dilation response amplitude (PDR), heart period response amplitude (HPR), top 10% *β*-estimate in the left dorsal horn of C6 (ROI), in the dilated left dorsal quadrant of C6 (dilated ROI), in the right ventral horn of C6 (control), and in the dilated right ventral quadrant of C6 (dilated control). Colors indicate ICC interpretation according to Cicchetti (1994): dark red: ICC < 0.4, poor; medium red: ICC 0.4 - 0.59, fair; orange: ICC 0.6 - 0.74, good; yellow: ICC 0.75 - 1.0, excellent. Individual values underlying the ICC calculation are shown in Supplementary Figure 1.

**Table 1.**
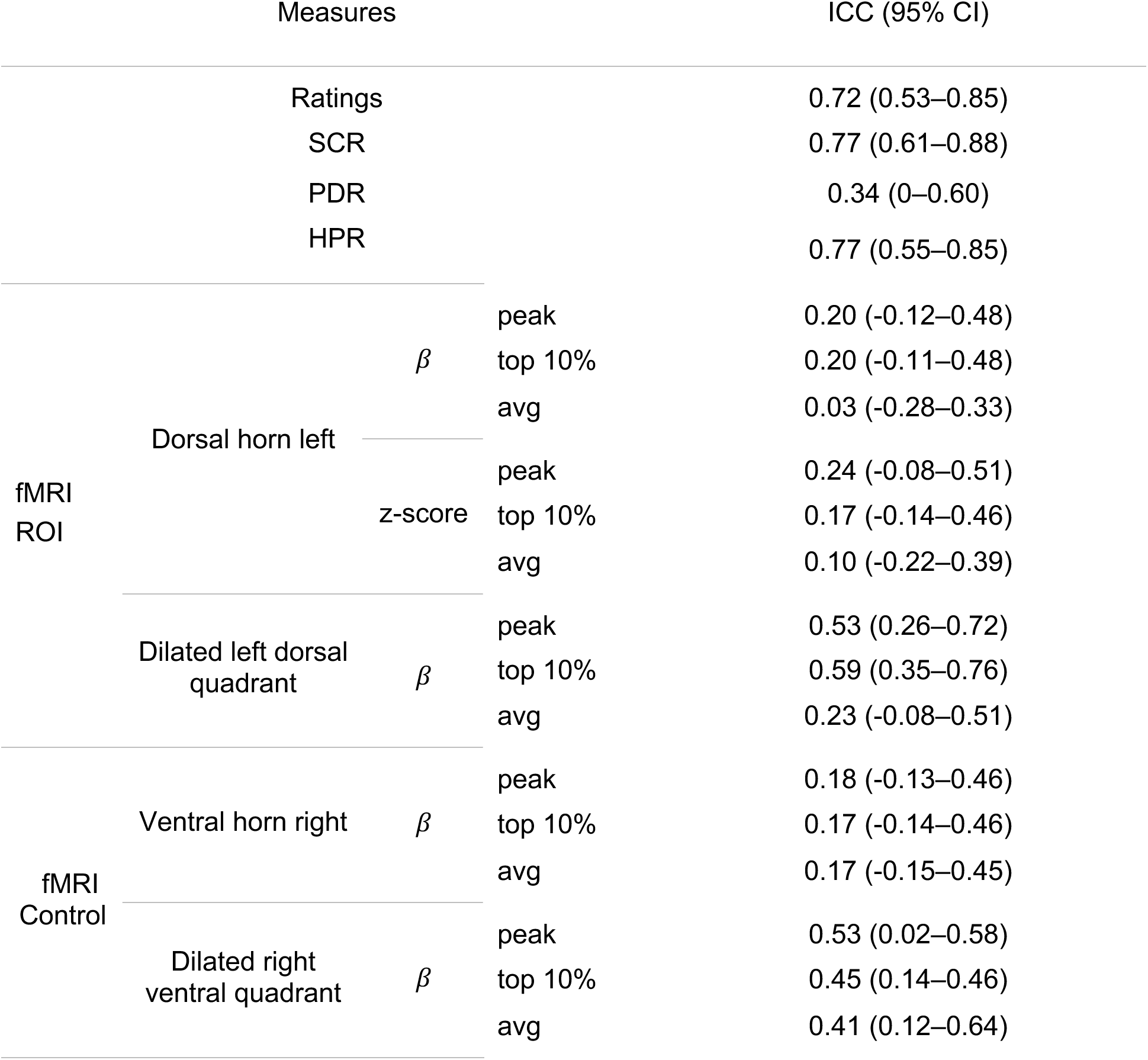
Intraclass correlation coefficient (ICC) and 95% confidence interval for subjective ratings, peripheral physiological data and BOLD response amplitudes.

#### 3.4.3 fMRI test-retest reliability

We calculated the reliability of the BOLD response amplitudes using four different masks: i) the left dorsal horn of C6 (ROI; grey matter area of interest), ii) an enlarged mask of the dorsal left cord quadrant of C6 (dilated ROI; including the venous plexus containing veins draining the dorsal horn), iii) the right ventral horn of C6 (control; grey matter area of no interest), and iv) an enlarged mask of the right ventral quadrant of C6 (dilated control; including the venous plexus with veins draining the ventral horn). To our surprise, all investigated metrics (*β* estimates of i) peak voxel, ii) top 10% of voxels, and iii) average across all voxels) showed poor reliability (all ICC < 0.4) in our target region, i.e., the left dorsal horn. This pattern did not change when we also took into account noise at the individual level, by repeating these analyses on z-values instead of *β*-estimates (with the latter only reflecting response amplitudes without taking into account residual noise; see Table 1). When assessing a larger region however (including draining vein territory), reliability was in the poor to fair range (ICC between 0.23 and 0.59). For the control areas, reliability in the gray matter region was consistently in the poor range (all ICC < 0.4), and in the poor to fair range (ICC between 0.41 and 0.53) in the dilated control region (as shown in Fig. 7).

#### 3.4.4 fMRI spatial consistency

To also compare the spatial patterns of the BOLD responses between days, we calculated Dice coefficients (DC). On the group level, the p-maps in the left dorsal horn of C6 (at p < 0.05 corrected) did not show any overlap across days (DC = 0). Using a more liberal thresholding increased the DC slightly (uncorrected p < 0.01: DC = 0.009, uncorrected p < 0.05: DC = 0.26). On the individual level (using an uncorrected p < 0.05), z-maps of 35 participants held supra-threshold voxels in the left dorsal horn (C6), but in only 5 participants these overlapped across days (mean DC across all 35 participants with suprathreshold voxels in ROI on both days: 0.05, mean DC of 5 participants with overlap: 0.33, range 0.02 – 0.65).

### 3.5 Post-hoc analyses

#### 3.5.1 Increased number of runs

In order to assess whether an increase in stimulus numbers would lead to a higher reliability, we used all runs with a TR of 1800ms (four runs per day instead of just one). When first assessing response amplitude, we noticed that this led to a clear enhancement in the strength of the group-level BOLD response, not only in the average across days (t(39) = 6.64, p_corr_ < 0.001, 331 supra-threshold voxels), but also for Day 1 (t(39) = 4.77, p_corr_ = 0.002, 94 supra-threshold voxels) and for Day 2 separately (t(39) = 5.55, p_corr_ < 0.001, 285 supra-threshold voxels; Supplementary Figure 2). Most importantly, with the increased stimulus numbers we now observed an overlap of activation across both days (94 supra-threshold voxels at p < 0.05 corrected), leading to an improved dice coefficient (DC = 0.80 for group-level p-maps at p_corr_ < 0.05), meaning that spatial consistency improved substantially. Contrary to our expectations, increasing the number of trials did not lead to improvements in the reliability of the non-BOLD data or the BOLD response amplitude in the target region (Supplementary Fig. 3 and Supplementary Table 2).

#### 3.5.2 Accounting for spontaneous activity

When accounting for spontaneous fluctuations of BOLD activity in the left dorsal horn by adding the time-course of the right dorsal horn as a slice-wise regressor to the GLM, we did not observe an increase in test-retest reliability (Supplementary Fig. 3 and Supplementary Table 2).

#### 3.5.3 Within-run reliability

Interestingly, the within-run reliability depended on the selection of trials. Between odd and even trials, reliability in the target region (left dorsal horn, C6) reached the level “good” (ICC = 0.69, Supplementary Fig. 3 and Supplementary Table 2) for the top 10% *β* estimates, and still “fair” for the ROI average (ICC = 0.52), whereas comparing the first half of trials to the second resulted in poor reliability (ICC = 0.36, top 10% *β*). Odd-even reliability was excellent for all three peripheral measures (Supplementary Fig. 3 and Supplementary Table 2), and slightly lower, but still fair to excellent, for the first vs. second half of trials.

## 4. Discussion

In this study we aimed to probe the limitations of task-based spinal fMRI – a young field facing many challenges – by investigating the robustness of spinal cord BOLD responses to repeated nociceptive stimulation across two consecutive days. For this purpose, we first examined if BOLD activation patterns occurred in the expected region of the spinal cord and assessed the spatial specificity of the response across a larger area. In our main investigation, we focused on the test-retest reliability of the BOLD response amplitude as well as the consistency of its spatial pattern. To disambiguate between effects on reliability of either data quality or variability in the underlying process of nociception, we also assessed the reliability of several simultaneously-recorded peripheral physiological measures.

### 4.1 Heat-pain evoked responses

In order to ascertain that the chosen stimulation parameters (contact heat stimuli at 48°C for 1s) would elicit a robust response, we recorded subjective ratings as well as peripheral physiological responses. Our results are in line with the general observation that painful stimulation activates the autonomic nervous system (Boucsein, 2012; Cowen et al., 2015; Kyle & McNeil, 2014), exemplified here by increased skin conductance, pupil dilation and heart rate in response to the stimulus. Although these responses are not specific for pain per se, as they can generally indicate increased arousal or salience (Lee et al., 2020), along with perceptual ratings being clearly above pain threshold, they suggest that a robust pain response was evoked by our brief nociceptive stimulation.

Our data furthermore showed that a brief contact heat stimulus of 1s can already evoke a measurable group-level BOLD response in the dorsal horn of the spinal cord. This response was observed in the expected segmental level and survived strict permutation-based correction for multiple comparisons. In humans, there is ample evidence that heat pain stimulation leads to activation in the ipsilateral DH of the spinal gray matter at the expected rostrocaudal location (Bosma & Stroman, 2015; Brooks et al., 2012; Eippert et al., 2009; Geuter & Buchel, 2013; Nash et al., 2013; Oliva et al., 2022; Seifert et al., 2023; Sprenger, Eichler, et al., 2018; Summers et al., 2010; Weber et al., 2016b, see Kolesar et al., 2015 for review), though these studies have consistently used longer stimulus durations. While results from one spinal fMRI study in the motor domain suggested that short stimuli may elicit weaker BOLD responses than expected (Giulietti et al., 2008), the application of short stimuli in event-related designs allows for a larger number of trials and may therefore boost power, as well as enable greater variability in the timing of the stimulus presentation (D’Esposito et al., 1999). Such features could be helpful for investigating the cognitive modulation of pain (Atlas & Wager, 2012; Villemure & Bushnell, 2002; Wiech, 2016) at the spinal level with more sophisticated paradigms than currently employed.

### 4.2 Spatial specificity

After confirming that BOLD responses indeed occurred in the ipsilateral dorsal horn of the expected segment, we next investigated the response pattern beyond this target area, to allow insights into the spatial specificity of BOLD responses. In the target segment of the spinal cord (C6), the ipsilateral dorsal horn indeed showed the highest number of active voxels across all four gray matter horns, though in all horns smaller numbers of active voxels were observed. Activation beyond the ipsilateral dorsal horn has been reported in previous spinal cord fMRI studies (Cahill & Stroman, 2011; Geuter & Buchel, 2013; Summers et al., 2010; Yang et al., 2015; see Kolesar et al., 2015 for review) and one factor contributing to this could be spatial inaccuracies, e.g. due to distortions during image acquisition, suboptimal registration to template space and spatial smoothing (Bosma & Stroman, 2015; Eippert, Kong, Jenkinson, et al., 2017; Hoggarth et al., 2022). On the other hand, activations in the ventral horns as well as the contralateral dorsal horn could also be indicative of neural processing in these areas: not only is there evidence for functional connectivity between the dorsal horns in the human spinal cord (for review, Harrison et al., 2021), but also evidence from animal models for dorsal commissural interneurons (Bannatyne, 2006; Petkó & Antal, 2000) and primary afferents that project contralaterally (Culberson et al., 1979; Light & Perl, 1979). Furthermore, autoradiographic rat data also show widespread responses to noxious heat stimuli (Coghill et al., 1991, 1993) and a painful heat stimulus might trigger motor responses as well as the active inhibition of thereof (Pierrot-Deseilligny & Burke, 2012; Purves et al., 2019). Taking these aspects into account, we would argue that activations outside of the ipsilateral dorsal horn likely reflect more than noise.

Apart from activation in spinal cord segment C6, we also observed active voxels in segments C5, C7 and C8 (when using a liberal uncorrected threshold), a pattern that has been observed previously in human data (Geuter & Buchel, 2013; Rempe et al., 2015; Seifert et al., 2023; Weber et al., 2016a). In addition, Shekhtmeyster et al. (2023) provide evidence for cross-segmental spinal cord activation of glial cells of mice in response to nociceptive stimulation, hinting at similarly widespread spinal processing mechanisms. On the one hand, a part of this large rostrocaudal extent could be due to dermatomal variability between participants and the fact that adjacent spinal roots can innervate overlapping areas of skin (Lee et al., 2008). However, this aspect seems to be more relevant for tactile as opposed to pain and temperature dermatomes (Lorenz et al., 1996; Sherrington, 1898, as cited in Lee et al., 2008) and may therefore not explain the activity patterns we observed. Interestingly however, it has also been suggested that the size of dermatomes depends on central communication via spinal levels (Denny-Brown et al., 1973; Denny-Brown & Kirk, 1968; Kirk & Denny-Brown, 1970), emphasizing a dynamic view of cutaneous innervation. Building on this, the large rostrocaudal patterns may also be explained by inter-segmental nociceptive processing via e.g. propriospinal neurons (Flynn et al., 2011; Pierrot-Deseilligny & Burke, 2012) or inter-segmental projection patterns of primary afferents (Kato et al., 2004; Pinto et al., 2010).

Beyond the gray matter of the dorsal horn, the activation seemed to bleed into the subarachnoid space, with strong peaks just outside of the spinal cord, where the large veins that drain the spinal cord are located (Thron, 2016). Typically, the evaluation of BOLD effects in the spinal cord is restricted to either a cord mask or a gray matter mask, meaning that the extent to which such activation patterns are prevalent in the literature is uncertain. However, spinal fMRI studies employing a hypercapnia challenge reported stronger signal changes at the edge of the cord and in the CSF compared to inside the spinal cord (Barry et al., 2021; Cohen-Adad et al., 2010) and several spinal fMRI studies employing painful heat stimuli also reported activations on the outer edge of the cord or even overlapping into the CSF (Geuter & Buchel, 2013; Nash et al., 2013; Oliva et al., 2022; Rempe et al., 2015; Sprenger et al., 2015; Sprenger, Stenmans, et al., 2018). The bias towards draining veins is a well-known drawback of gradient-echo EPI and – considering our results – it might be advisable in the future to try disentangling micro- and macrovascular contributions to the spinal cord BOLD response, for instance by modeling the respective time courses (Kay et al., 2020), leveraging differences in TE- (Markuerkiaga et al., 2021; Uludağ et al., 2009) and phase dependencies (Stanley et al., 2021) to remove signal contributions from large veins, or suppressing draining vein signal during data acquisition (Li et al., 2022). A first proof-of-principle step might however be to obtain individual vasculature maps and investigate the relationship between vascular anatomy and BOLD activation patterns in the spinal cord, a concept that aligns with the initial approach of Cohen-Adad et al. (2010), whose findings highlight the importance of vascular dynamics.

### 4.3 Test-retest reliability across consecutive days

The main objective of this study was to investigate the test-retest reliability of task-based spinal cord BOLD responses across two consecutive days. Test-retest reliability describes to what extent repeated measurements yield similar results, given that the underlying true value has not changed (Lavrakas, 2008) and this also applies to inter-individual variation, with consistent differences between participants indicating good reliability. Considering that many factors contribute to the processing of pain (Bushnell et al., 2013; Heinricher & Fields, 2013), we also collected verbal ratings as well as peripheral physiological responses in response to painful stimulation – apart from offering a different window on the reliability of pain processing, these measures also served as controls against which we could compare the reliability of the BOLD responses. We observed that verbal ratings exhibited high reliability, a finding that has also been reported in previous studies (Letzen, 2014; Quiton & Greenspan, 2008; Upadhyay, 2015), though it is unclear to what extent this reflects the stability of actual perceptual differences or might be driven by biases in reporting pain or differences in interpreting the rating scale. Peripheral physiological measures also mostly showed high reliability, providing complementary evidence that participants can indeed be distinguished reliably based on their peripheral physiological response to pain (the low reliability of pupil dilation is likely due to noisy data on account of the non-ideal setup for eye-tracking with the 64-channel coil employed here). Together these data provide a solid foundation for investigating the reliability of spinal cord BOLD responses, as they indicate a generally high reliability of supra-spinal measures of pain processing.

When looking at the test-retest reliability of spinal cord BOLD responses across days, we observed that the reliability of the response amplitudes in the region of interest (left dorsal horn of segment C6) was consistently in the poor range, similar to results obtained by Weber and colleagues when investigating within-day test-retest reliability of spinal cord BOLD responses to heat-pain stimulation (Weber et al., 2016a). One could argue that two of our chosen metrics (ROI mean and peak) are suboptimal for assessing reliability, as the former included many non-responsive voxels and the latter may merely constitute an outlier (given the tSNR of the data). However, the more constrained approach of using the top 10% resulted in very similar reliability and higher reliability values have been reported when investigating BOLD responses to painful stimulation using similar approaches in the brain (Bi, 2021; Gay et al., 2015; Letzen, 2014; Quiton et al., 2014; Upadhyay, 2015). While the majority of these studies used stimuli longer than 10s (Gay et al., 2015; Quiton et al., 2014; Upadhyay, 2015), Letzen et al. (2014) reported good test-retest reliability for 4s contact heat stimuli and lower trial as well as participant numbers, similarly to Bi et al. (2021), who observed fair to moderate test-retest reliability when using 4ms radiant heat-laser stimuli on roughly half the number of participants compared to our study, albeit with slightly higher trial numbers. Partly, these differences may be explained by the larger tSNR typically achieved in brain compared to spinal cord fMRI as well as the lower spatial resolution employed in these studies, which further increases tSNR.

Interestingly, an extended mask that covered the venous plexus surrounding the spinal cord yielded good reliability for the top 10% of the parameter estimates. One possible interpretation of this finding is that spinal cord BOLD response amplitudes could indeed be a reliable measure, but with the employed gradient-echo EPI acquisition’s sensitivity to macrovascular responses (Bandettini et al., 1994; Duong et al., 2003; Gati et al., 1997; Uludağ et al., 2009), the actual response peak might be shifted from the gray matter towards the draining veins – in brain fMRI such differences would not be immediately noticeable, considering the typically lower spatial resolution, potentially causing signals from veins and gray matter to blend within individual voxels. Furthermore, the brain’s anatomical structure, where large draining vessels often lie directly on top of the cortical gray matter, contrasts with the spinal cord’s architecture, in which these larger draining vessels are located outside the white matter, surrounding the cord (Duvernoy, 1999; Gray, 2021).

In addition to response amplitudes, we also investigated the spatial consistency of the response pattern, i.e., if active voxels overlapped across days. The group level results showed a significant response in the target region for each session separately, but, the activation patterns did not overlap between days – while the location on the dorsal-ventral dimension remained similar, the patterns differed rostro-caudally within the same segment. This was paralleled on the individual level, where only few participants had overlapping responses in the area of interest, leading to very low dice coefficients at either level. It is noteworthy that a higher-than-expected amount of spatial variability in the z-direction within participants upon thermal stimulation has also been reported in a recent within-day design by Seifert and colleagues (2023). It is currently unclear if this is an indicator of large variability in spinal nociceptive processing, or the result of low tSNR (due to residual noise), an insufficient number of trials or a low stimulus intensity (Upadhyay, 2015). A partial answer was given by a post-hoc analysis where we increased the amount of data by averaging over multiple runs per session: here, the spatial overlap on the group level improved substantially, yet the reliability of the response amplitude did not improve.

There are several factors specific to across-day set-ups that could have negatively impacted the reliability of both response amplitudes and spatial patterns. One such factor is the quality of the spatial normalization, since differences thereof between days could have adverse effects on reliability (especially for a small structure such as the DH), yet this did not receive support by our analyses based on EPI-template Dice coefficients. Further inconsistencies between sessions may have been caused for example by differences in the positioning of the participants in the scanner, resulting in a different tilt of the head and neck, which is supported by the moderate correlation between differences in BOLD parameter estimates and the angulation of the slice stack relative to the static magnetic field (as a proxy for neck positioning). Due to the anatomical organization of the draining veins, the resulting different curvature / orientation of the neck and thus spinal cord across days may have impacted the relative contribution of the longitudinal and radial veins to the overall signal (Giove et al., 2004; Viessmann et al., 2019). It is also possible that the general physiological state of the participants produced some variation the fMRI responses (Dubois & Adolphs, 2016), but except for a slight increase in heart rate from day one to day two, all other markers of physiological arousal remained the same. In one post-hoc analysis, we tried to partially address these limitations by computing the within-run reliability using odd/even trials as well as the first vs. second half of trials, where no re-positioning in the scanner occurred and spatial normalization was equal. Interestingly, odd-even reliability of the BOLD fMRI results was in the good range, while early-late reliability was still poor. The main difference between these two assessments is the temporal distance between the trials, which was greater for early-late reliability. This indicates that the heat-pain evoked activations display considerable variability within a single run, potentially due to mechanisms physiological mechanisms such as adaptation, habituation and sensitization (Greffrath et al., 2007; Hollins et al., 2011; Latremoliere & Woolf, 2009) as well as potential technical issues such as scanner drift. While it is unlikely that the same mechanisms account for across-day variability, it indicates that factors beyond positioning and spatial normalization can contribute to low reliability.

It is important to point out that a systematic comparison of our results with reliability estimates obtained by resting-state spinal cord fMRI studies is unfortunately not possible, as these studies not only varied vastly in their sample sizes (N = 1 to N = 45) but consistently used a within-day design (Barry et al., 2016; Hu et al., 2018; Kaptan et al., 2023; Kong et al., 2014; Liu et al., 2016; San Emeterio Nateras et al., 2016), thus not encountering the issues of across-day measurements that might bring about low reliability. A notable exception is a recent study by Kowalczyk and colleagues (2024), where a between-day design also resulted in mostly ‘poor’ voxelwise ICCs, though the spatial patterns of connectivity showed near-perfect agreement.

### 4.4 General considerations on reliability

To put our observation of mostly low reliability of spinal cord BOLD responses into a larger context, it is important to mention that a recent meta-analysis investigating univariate BOLD responses in the brain to several common tasks from various domains also observed generally low reliability (Elliott et al., 2020). Several ways to improve the reliability of fMRI have been discussed (Elliott et al., 2021; Kragel et al., 2021), such as multivariate analysis (Gianaros et al., 2020; Han et al., 2022), modeling stable variability, and the aggregation of more data (Elliott et al., 2021), all of which might be applicable in the context of spinal cord fMRI as well.

A further aspect deserving discussion is our quantification of reliability, which was carried out using the intra-class correlation coefficient ICC(3,1) (Shrout & Fleiss, 1979), a common measure for test-retest reliability of fMRI data (Caceres et al., 2009; Elliott et al., 2020; Noble et al., 2021). The ICC is a useful metric to investigate inter-individual differences, since it quantifies to what extent participants can be “re-identified” across repeated measurements by means of the stability of their rating in relation to that of other participants (Brandmaier et al., 2018; Hedge et al., 2018; Liljequist et al., 2019). In order to obtain a high ICC, the variation between participants should be large, and the variation within participants as well as the general measurement error should be small. However, traditional univariate analyses of BOLD responses via the GLM – as also employed here – are designed to minimize between-participant variability in order to gain a robust group-level response (Fröhner et al., 2019; Hedge et al., 2018). Given the possible sources of noise discussed in this study and elsewhere (Eippert, Kong, Jenkinson, et al., 2017; Kinany, Pirondini, Micera, et al., 2022; Summers et al., 2014), minimizing the measurement noise holds the potential to both improve reliability and optimize main effects on the group level.

### 4.5 Limitations

There are several limitations of this work that need to be considered. First, the BOLD responses elicited by 1s contact heat stimuli may exhibit lower reliability compared to those from longer stimulus durations in a block design, which are typically more effective in detecting effects and could yield more robust activation patterns (Bennett & Miller, 2013). The limited number of trials further constrains our assessment of test-retest reliability, potentially making it more restrictive than studies using more powerful experimental designs; here it is also important to mention the low degrees of freedom of our time-series (considering that only 160 volumes were acquired per run and that extensive denoising was carried out). Second, an assessment of the test-retest reliability of tSNR values – as possible for example via a short resting-state acquisition – in the left dorsal horn across different days would have provided valuable insights into the consistency of the fMRI signal quality and should be considered in future studies. Third, one might argue that instead of delivering stimuli with the same temperature on both days, we could have instead matched stimulus intensity across days based on subjectively perceived intensity (to account for confounds that might differ across days). Fourth, while the use of MP-PCA resulted in a substantial tSNR increase (without a strong spatial smoothness penalty), future studies might look in more detail at potential violations of underlying assumptions as well as artificial activation spreading, which has recently been observed under certain conditions (Fernandes et al., 2023). Finally, we might have achieved an increased across-day reliability by minimizing variability in participant position (and thus also spatial inaccuracies), for example by using personalized casts (Power et al., 2019).

### 4.6 Conclusion

We observed that heat pain stimuli as short as 1s can evoke a robust BOLD response in the ipsilateral dorsal horn of the relevant spinal cord segment, making such stimuli suitable for use in cognitive neuroscience experiments that require variable trial designs and large numbers of trials. Although autonomic and subjective indicators of pain processing showed mostly good-to-excellent reliability, BOLD response patterns varied strongly within participants, resulting in poor test-retest reliability in the target region. Interestingly, using an extended analysis region including the draining veins improved reliability across days, suggesting that future studies should aim to disentangle macro- and microvascular contributions to the spatial response profile. Our results indicate that further improvements in data acquisition and analysis techniques are necessary before event-related spinal cord fMRI can be reliably employed in longitudinal designs or clinical settings. To facilitate such endeavours, all data and code of this study are publicly available, thus allowing others to develop and improve pre-processing and analysis strategies to overcome current limitations.

## Data and code availability

The underlying data and code are currently only accessible to reviewers, but will be made openly available upon publication via OpenNeuro and GitHub, respectively.

## Ethics

All participants gave written informed consent. The study was approved by the Ethics Committee of the Medical Faculty of the University of Leipzig.

## Author contributions

Author contributions are listed alphabetically according to CRediT taxonomy (https://credit.niso.org).

Conceptualization: AD, FE

Data curation: AD

Formal analysis: AD, FE, UH

Funding acquisition: FE

Investigation: AD

Methodology: AD, FE, JL, RM

Project administration: AD, FE

Resources: JF, JL, RM

Software: AD, FE, UH, MK

Supervision: FE, TM, NW

Visualization: AD

Writing – original draft: AD, FE

Writing – review & editing: JB, AD, FE, JF, UH, MK, JL, RM, TM, NW

## Acknowledgements

We would like to thank all volunteers who participated in this study. Additionally, we want to thank everyone who assisted in data acquisition as well as Lisa-Marie Pohle for her help in the randomization of trials and conditions. This manuscript will be part of a doctoral thesis.

## Supplementary Material

**Supplementary Figure 1.**
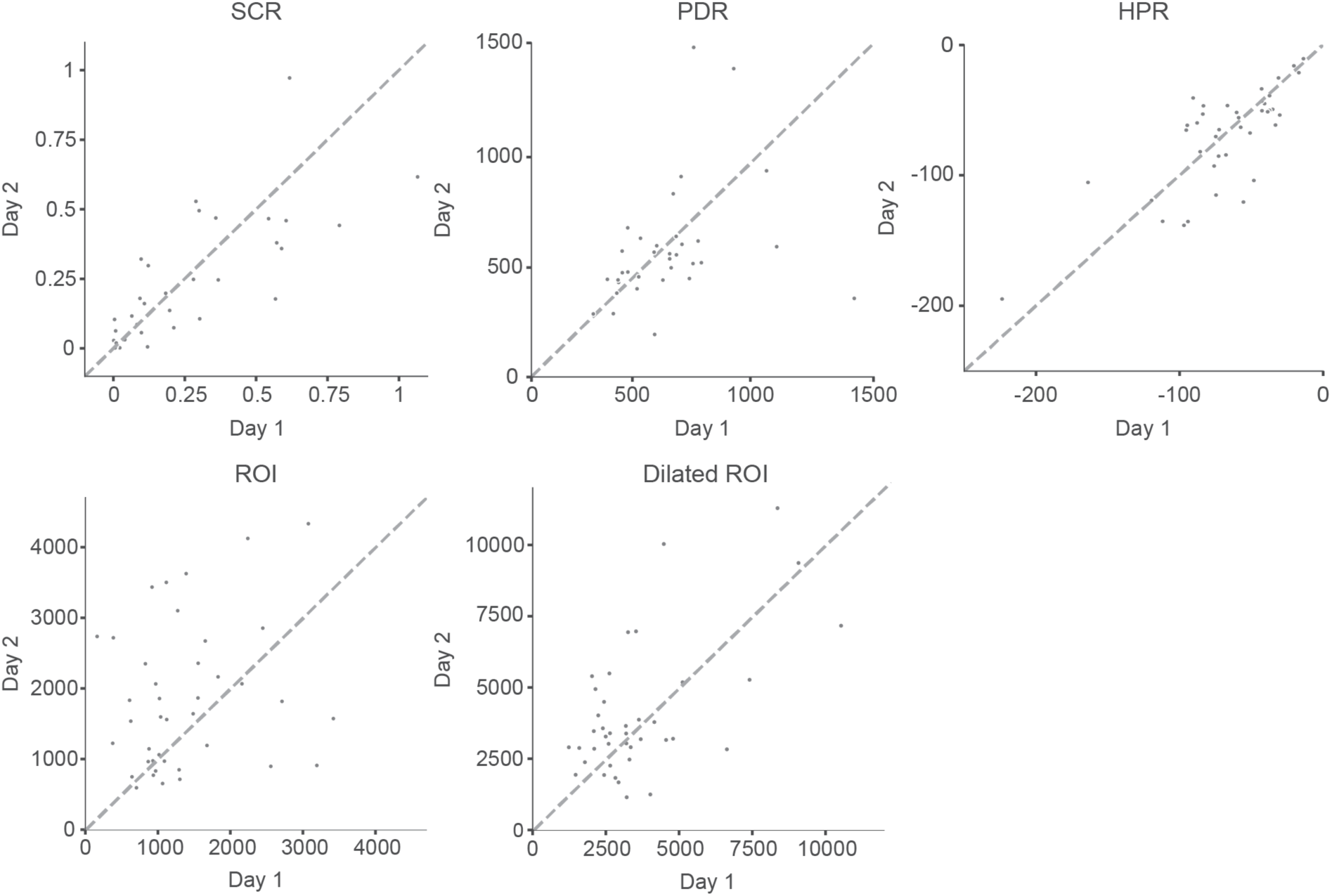
Individual values underlying ICC calculation. Participant-wise data of Day 1 and Day 2 peak values for skin conductance responses (SCR), pupil dilation responses (PDR) and heart period responses (HPR), as well as average top 10% *β* values of the left dorsal horn (ROI) and the dilated left dorsal quadrant (Dilated ROI) in spinal cord segment C6.

**Supplementary Figure 2.**
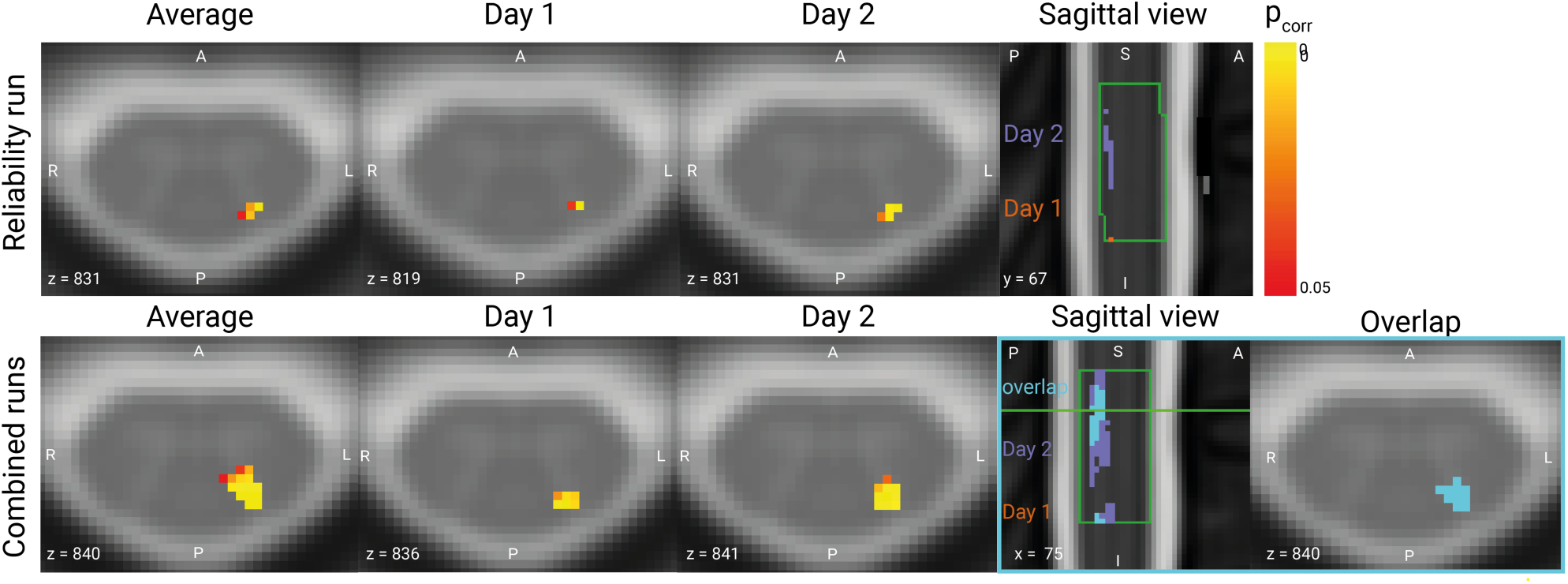
Group-level fMRI results. Results are shown for the Reliability Run (i.e., one run per day; top row; same image as in main manuscript used for comparison purposes here) and Combined Runs (i.e. average of four runs per day; bottom row). Only voxels surviving a threshold of p < 0.05 (corrected for multiple comparisons via a permutation test in a mask of the left dorsal horn in spinal cord segment C6) are displayed on top of a T2*-weighted spinal cord template (PAM50) in axial view, and on top of a T2-weighted spinal cord template (PAM50) in the sagittal view. In contrast to the Reliability Run, the Combined Runs demonstrated substantial overlap across both days. Spinal cord segment C6 is outlined in green in the sagittal images.

**Supplementary Figure 3.**
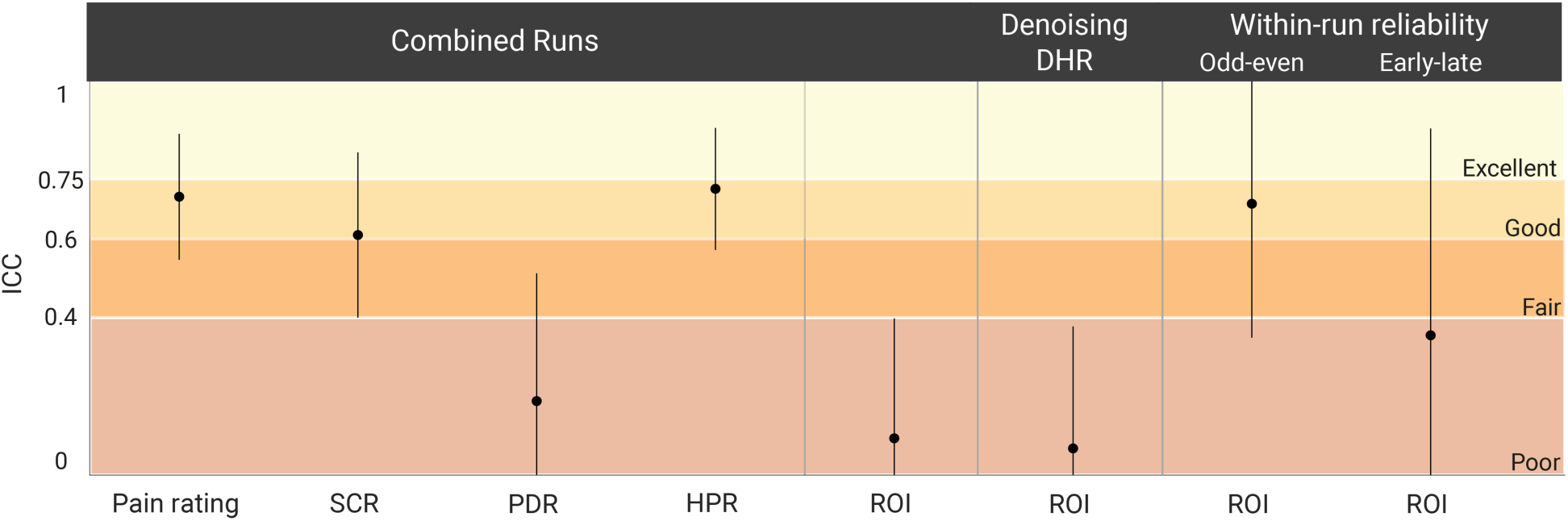
Test-retest reliability across both days for subjective ratings, peripheral physiological data and BOLD response amplitudes. Reliability is indicated via ICCs (plotted as dots with 95% CI represented as a line). *Combined Runs*: ICCs are reported for (from left to right) verbal ratings, SCR, PDR, HPR and top 10% *β*-estimate in the left dorsal horn of C6 (ROI). *Denoising DHR*: ICC of the top 10% *β*-estimate in the left dorsal horn of C6 (ROI) of only the Reliability Run, obtained from a GLM with an additional regressor of right dorsal horn activity. *Within-run reliability*: ICCs obtained from comparing either odd and even trials numbers (odd-even), or the first and second half of a run (early-late). Colors indicate ICC interpretation according to Cicchetti (1994): dark red: ICC < 0.4, poor; medium red: ICC 0.4 - 0.59, fair; orange: ICC 0.6 - 0.74, good; yellow: ICC 0.75 - 1.0, excellent.

**Supplementary Table 1.**
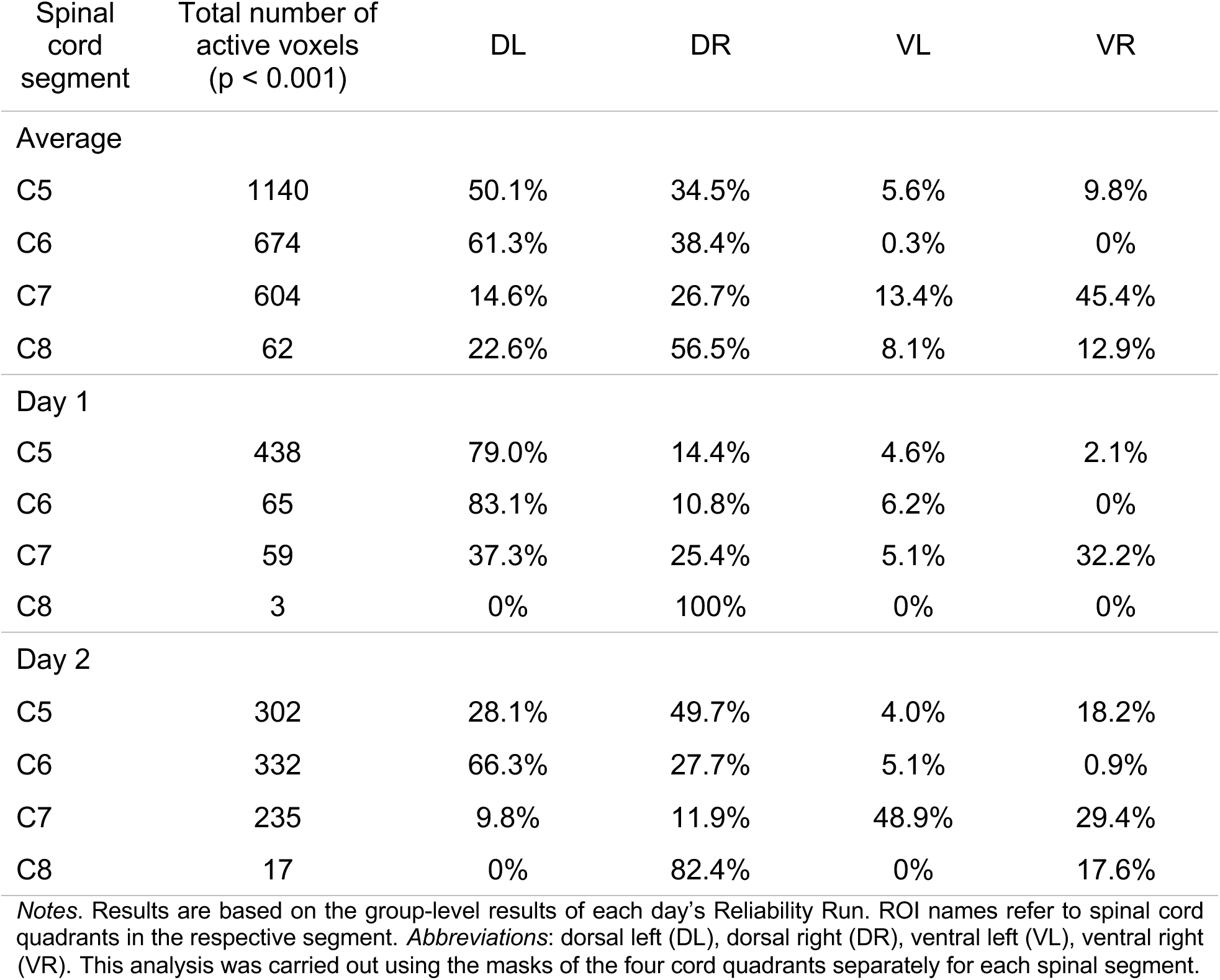
Percent suprathreshold voxels in the four cord quadrants of each spinal segment.

| Spinal cord segment | Total number of active voxels (p < 0.001) | DL | DR | VL | VR |
| --- | --- | --- | --- | --- | --- |
| Average |  |  |  |  |  |
| C5 | 1140 | 50.1% | 34.5% | 5.6% | 9.8% |
| C6 | 674 | 61.3% | 38.4% | 0.3% | 0% |
| C7 | 604 | 14.6% | 26.7% | 13.4% | 45.4% |
| C8 | 62 | 22.6% | 56.5% | 8.1% | 12.9% |
| Day 1 |  |  |  |  |  |
| C5 | 438 | 79.0% | 14.4% | 4.6% | 2.1% |
| C6 | 65 | 83.1% | 10.8% | 6.2% | 0% |
| C7 | 59 | 37.3% | 25.4% | 5.1% | 32.2% |
| C8 | 3 | 0% | 100% | 0% | 0% |
| Day 2 |  |  |  |  |  |
| C5 | 302 | 28.1% | 49.7% | 4.0% | 18.2% |
| C6 | 332 | 66.3% | 27.7% | 5.1% | 0.9% |
| C7 | 235 | 9.8% | 11.9% | 48.9% | 29.4% |
| C8 | 17 | 0% | 82.4% | 0% | 17.6% |

Notes. Results are based on the group-level results of each day's Reliability Run. ROI names refer to spinal cord quadrants in the respective segment. Abbreviations: dorsal left (DL), dorsal right (DR), ventral left (VL), ventral right (VR). This analysis was carried out using the masks of the four cord quadrants separately for each spinal segment.

**Supplementary Table 2.**
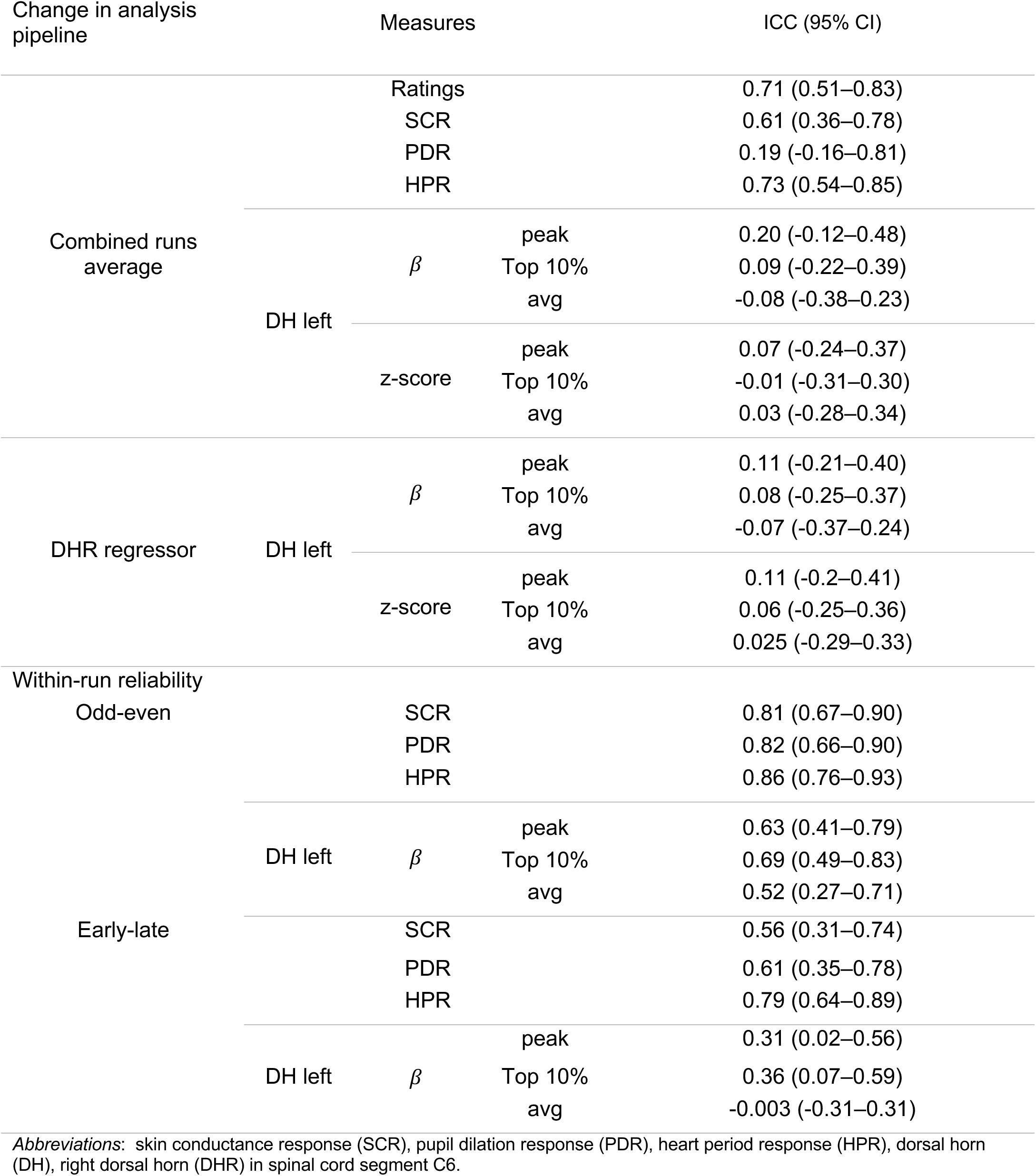
Intraclass correlation coefficient and 95% confidence interval for subjective ratings, peripheral physiological data and BOLD response amplitudes of post-hoc analyses.

**Supplementary Table 3.** Correlations between response measures.

| Peripheral /<br>subjective<br>measure | BOLD parameter estimate<br>(top 10%) | Pearson's r | p-value |
| --- | --- | --- | --- |
|  |  |  | One-tailed |
| Ratings | $\beta$ | -0.18 | 0.855 |
|  | z-score | -0.30 | 0.965 |
| SCR | $\beta$ | 0.34 | 0.017 * |
|  | z-score | 0.36 | 0.014 * |
| HPR | $\beta$ | -0.28 | 0.038 * |
|  | z-score | -0.28 | 0.039 * |
| PDR | $\beta$ | -0.11 | 0.722 |
|  | z-score | -0.05 | 0.611 |

*Notes.* Results are based on individual BOLD responses and peripheral physiological responses and subjective ratings of the reliability run. We correlated responses averaged across days for each measure. BOLD responses were quantified as the top 10 % of $\beta$ estimates and z-scores extracted from the left dorsal horn in spinal cord segment C6. *Abbreviations:* skin conductance response (SCR), pupil dilation response (PDR), heart period response (HPR). For further information see section 2.8.4. \* $p < 0.05$

**Supplementary Table 4.**
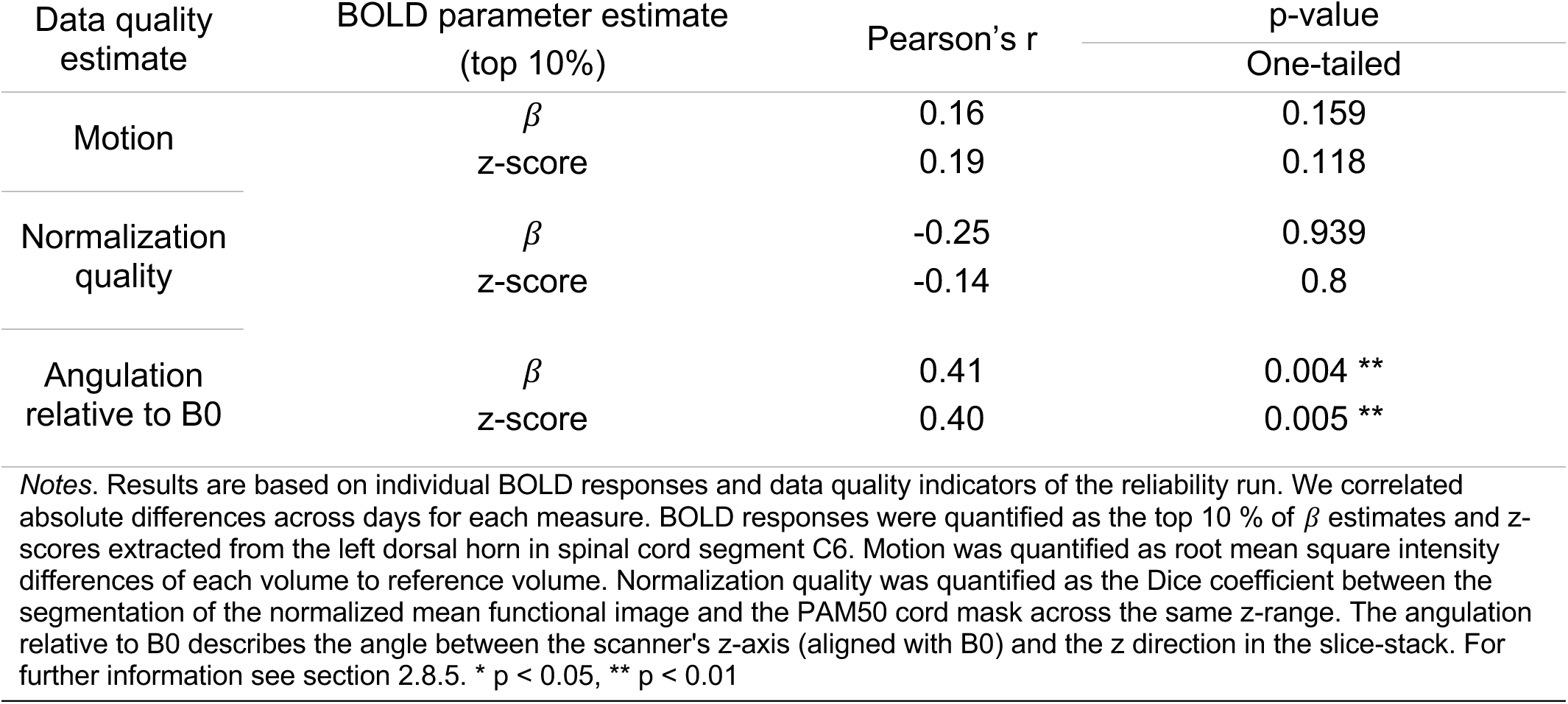
Correlations between BOLD parameter estimates and indicators of data quality.

| Data quality<br>estimate | BOLD parameter estimate<br>(top 10%) | Pearson's r | p-value |
| --- | --- | --- | --- |
|  |  |  | One-tailed |
| Motion | $\beta$ | 0.16 | 0.159 |
|  | z-score | 0.19 | 0.118 |
| Normalization<br>quality | $\beta$ | -0.25 | 0.939 |
|  | z-score | -0.14 | 0.8 |
| Angulation<br>relative to B0 | $\beta$ | 0.41 | 0.004 ** |
|  | z-score | 0.40 | 0.005 ** |

*Notes.* Results are based on individual BOLD responses and data quality indicators of the reliability run. We correlated absolute differences across days for each measure. BOLD responses were quantified as the top 10 % of $\beta$ estimates and z-scores extracted from the left dorsal horn in spinal cord segment C6. Motion was quantified as root mean square intensity differences of each volume to reference volume. Normalization quality was quantified as the Dice coefficient between the segmentation of the normalized mean functional image and the PAM50 cord mask across the same z-range. The angulation relative to B0 describes the angle between the scanner's z-axis (aligned with B0) and the z direction in the slice-stack. For further information see section 2.8.5. \* $p < 0.05$ , \*\* $p < 0.01$

## Deviations from preregistration

In the preregistration, we stated that in addition to ICC(3,1) we would report the Pearson correlation coefficient and ICC(2,1) as indicators of reliability. However, for the sake of brevity we ultimately decided to report only ICC(3,1), as all indicators were found to be highly similar.

In the preregistration, we stated that we aimed to calculate voxel-wise ICC maps. However, given that we observed almost no overlap of activation across days in the analysis reported here (thus making a voxel-wise assessment pointless), we decided to focus solely on the ROI assessments.

In the preregistration, we stated that we would investigate spatial aspects of reliability via x-, y-, and z-coordinates. However, for the sake of brevity we decided to instead employ Dice coefficients (i.e. a measure of spatial overlap), as we deemed them a more succinct and comprehensive representation of our data, also considering the complexity of the manuscript.

In the preregistration, we stated that we would assess reliability only in the ipsilateral dorsal horn of spinal cord segment C6. However, after observing robust activation extending to areas outside the gray matter, we chose to also investigate a larger region encompassing the draining vein territory.

Any other analyses carried out here, but not included in the preregistration, are clearly indicated as post-hoc analyses in the manuscript (see section 2.8).

